# Hippocampal engrams generate flexible behavioral responses and brain-wide network states

**DOI:** 10.1101/2023.02.24.529744

**Authors:** Kaitlyn E. Dorst, Ryan A. Senne, Anh H. Diep, Antje R. de Boer, Rebecca L. Suthard, Heloise Leblanc, Evan A. Ruesch, Sara Skelton, Olivia P. McKissick, John H. Bladon, Steve Ramirez

**Author notes:** These authors contributed equally to this work (Kaitlyn E. Dorst, Ryan A. Senne).

## Abstract

Memory engrams are both necessary and sufficient to mediate behavioral outputs. Defensive behaviors such as freezing and avoidance are commonly examined during hippocampal-mediated fear engram reactivation, yet how reactivation of these cellular populations across different contexts engages the brain to produce a variety of defensive behaviors is relatively unclear. To address this, we first optogenetically reactivated a tagged fear engram in the dentate gyrus (DG) subregion of the hippocampus across three distinct contexts. We found that there were differential amounts of light-induced freezing depending on the size of the context in which reactivation occurred: mice demonstrated robust light-induced freezing in the most spatially restricted of the three contexts but not in the largest. We then utilized graph theoretical analyses to identify brain-wide alterations in cFos co-activation during engram reactivation across the smallest and largest contexts. Our manipulations conferred greater positive cFos correlations and recruited regions spanning putative fear and defense systems as hubs in the respective networks. Moreover, reactivating DG-mediated engrams generated network topologies across experimental conditions, emphasizing both shared and distinct features. By identifying and manipulating the circuits supporting memory function, as well as their corresponding brain-wide activity patterns, it is thereby possible to resolve systems-level biological mechanisms mediating memory’s capacity to modulate behavioral states.

**SIGNIFICANCE STATEMENT:** Implementing appropriate defensive behaviors across disparate environments is essential for survival. Memories can be used to select these responses. Recent work identified and artificially manipulated cellular ensembles within the hippocampus that mediate fear memory recall, yet how these populations engage brain-wide pathways that mediate defensive behaviors under environmental contingencies is unclear. We demonstrated here that reactivation across environments of various sizes elicits different behavioral responses and corresponding brain-wide network dynamics. These findings establish the flexibility of memory-bearing ensembles in generating brain and behavior states.

## INTRODUCTION

All animals utilize a repertoire of defensive strategies to avoid danger under a variety of environmental conditions. For instance, it could be more advantageous to hide when hunted in a densely forested area, yet fleeing could be more advantageous when hunted in a vast field. The brain performs a series of computations to integrate important contextual information to then dictate appropriate behavioral outputs (Fanselow, 1994; Gross and Canteras, 2012; Silva et al., 2016). The defensive strategies chosen, such as freezing in a forest or fleeing in a field, can be learned and used to guide the animal in future scenarios.

Memory systems play an important role in mediating defensive actions based on past events. This helps the animal avoid potentially harmful scenarios or cope with similar threatening situations. The hippocampus is an evolutionary conserved region central to episodic memory processes that guide defensive actions during fear memory recall, and lesions show that hippocampal disruption can impair such behavioral responses (Fanselow, 1994; Kim et al., 1993; Maren et al., 2013; Scoviille and Milner, 1957). Populations of cells within the hippocampus, often termed “engram” ensembles, are both necessary and sufficient to drive memory expression (Chen et al., 2019; Josselyn and Tonegawa, 2020; Liu et al., 2012). Activity-dependent and inducible tagging systems have allowed researchers to artificially manipulate engrams encoding fearful experiences to drive defensive actions (Reijmers et al., 2007). Many of these studies measured freezing behavior when a hippocampal fear engram is reactivated in a novel context (Liu et al., 2012; Roy et al., 2016; Ryan et al., 2015). However, recent reports show that hippocampal fear engrams can drive place aversion or anxiety-related avoidance-like responses as well (Chen et al., 2019; Ramirez et al., 2013; Redondo et al., 2014). This alludes to the possibility tested here that hippocampal fear engrams are flexible in their capacity to drive state-dependent alterations in behavioral outputs contingent on external demands placed on the animal. How these discrete populations can engage underlying neural systems to gate defensive behaviors is relatively unknown.

To gain mechanistic insight into memory-driven defensive behaviors, we leverage activity-dependent, inducible tagging strategies to optogenetically manipulate hippocampal fear engrams in freely behaving mice. We first tagged a hippocampal fear engram in the dentate gyrus subregion (DG) with blue light-sensitive channelrhodopsin-2 (ChR2) during contextual fear conditioning (CFC). Over subsequent days, mice were then subjected to optogenetic reactivation in a battery of environments of various sizes. Our findings show that the tagged hippocampal CFC engram is not fixed to a singular defensive response during optogenetic reactivation, as mice engaged in more defensive actions such as “freezing” when the hippocampal fear engram was reactivated in a small arena but not in a large area.

Moreover, a variety of neural circuits are implicated in mediating defensive actions. These areas span putative “fear” and “defense” systems for sensory detection, integration, and commanding behavior output (Fendt and Fanselow, 1999; Gross and Canteras, 2012; Silva et al., 2016; Tovote et al., 2015). Recent technological advancements in microscopy allow researchers to take an unbiased approach to examine interactions between these systems with mesoscale resolution (Dean et al., 2015; Kim et al., 2015; Park et al., 2019; Swaney et al., 2019; Yun et al., 2019). Using this approach, we next examined brain-wide pairwise correlations in endogenous cFos expression in mice that underwent optogenetic reactivation of the hippocampal CFC engram in either the smallest arena or the largest, as those environments promoted and discouraged light-induced freezing, respectively. We then utilized network analyses based in graph theory to first examine the topological nature of the functional interactions between brain areas and then identify mediator “hub” regions that are crucial to the resulting graphs. We found that groups which showed stereotypical freezing behavior during recall (natural or engram reactivation) had significantly greater average clustering coefficients, that the hippocampus showed increased functional connectivity with hypothalamic areas, and that there were shared hub regions mediating memory and behavior.

Together, our results show that hippocampal CFC engrams drive behavioral outputs that are contingent on environmental parameters such as size. These outputs uniquely engage brain-wide processes and point to numerous hub regions as sites for future perturbation studies. The flexibility of hippocampal CFC engrams underscores the dynamic nature of memory-guided behavior and offers a new dimension to intervening with disorders of the brain in which fear is a core component, such as PTSD and anxiety.

## RESULTS

### Environment size influences light-induced freezing

We tested the capacity of a hippocampal CFC engram to mediate behavioral changes contingent on external demands placed on the animal. Mice first underwent CFC during the tagging phase for activity-dependent labeling of the dedicated engram ensemble with ChR2-eYFP or -eYFP (**Figure 1A-B**). Over subsequent days, they were subjected to optogenetic reactivation of the hippocampal CFC engram while exploring environments of various sizes (i.e., “Small Box” [SB], “Mid Box” [MB], “Large Box” [LB]). These environments were contextually distinct from the original arena in which they experienced CFC to prevent generalization (**Figure 1C-D**; See Methods). We quantified freezing behavior, which is defined as the cessation of all movement with the exception of breathing (Grossen and Kelley, 1972), as this is a common rodent behavioral metric of a negative affective state such as fear.

**Figure 1:**
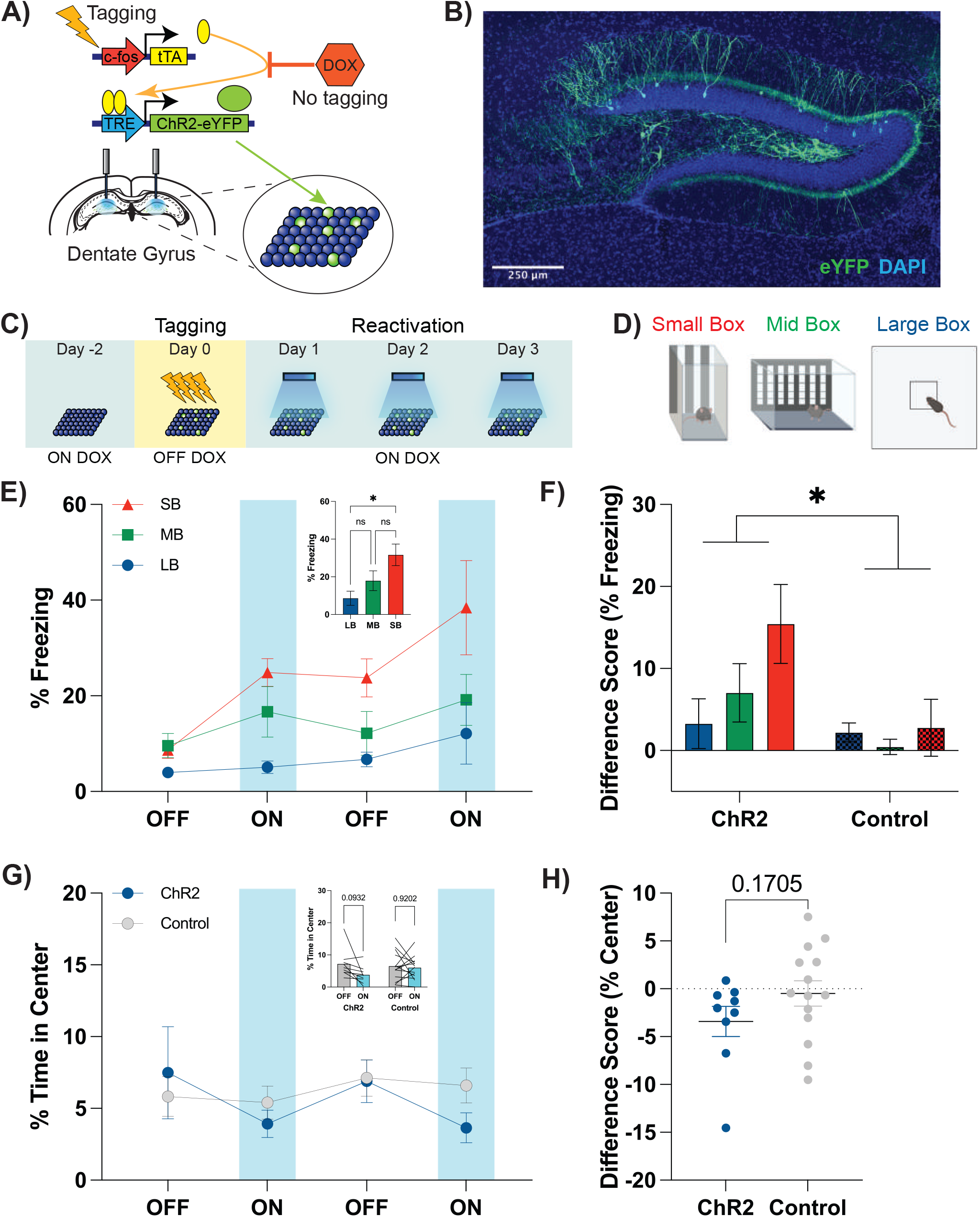
Engram reactivation drives light-induced freezing contingent on environment size. (A) Schematic of activity-dependent tagging of engram ensembles in the dentate gyrus (DG). (B) Representative image of a tagged Contextual Fear Conditioning (CFC) engram in the DG. Scale bar represents 250 μm. (C) Experimental design for hippocampal CFC engram tagging and reactivation. Mice (ChR2, n = 9; Control, n = 14) are taken off of doxycycline (DOX) two days prior to hippocampal CFC engram tagging. After a series of foot shocks are delivered during tagging (4 shocks, 0.5 mA, 2 seconds), all mice are placed back on DOX. The tagged hippocampal CFC engram is reactivated across three days. (D) Novel environments of various sizes used for hippocampal CFC engram reactivation (Small Box [SB], Mid Box [MB], Large Box [LB]). All mice are randomly assigned to an order of environments for reactivation in Figure 1C. (E) %Freezing levels for all ChR2 animals across light epochs. There is a significant increase in the amount of light-induced freezing in the SB compared to LB (inset). (F) Difference scores (Average %Freezing light-on - Average %Freezing light-off) generated from ChR2 and Control animals. There is a main effect of virus condition on the difference score, but no effect from environment size or an interaction between the two. (G) %Time in Center for ChR2 and Control animals in the LB across light epochs. There is no significant difference in the amount of time spent exploring the center of the chamber across groups. (H) Difference scores (Average %Time in Center light-on - Average %Time in Center light-off) generated from ChR2 and Control animals. There is no significant difference. Data are presented as Mean ± SEM. Significant differences are reported as (*) p < 0.05.

Strikingly, we observed that optogenetic reactivation of a hippocampal CFC engram was sufficient to induce freezing in the Small Box, whereas in the same animals light-dependent freezing behavior was not observed in the Large Box. Importantly, we found that there was no difference in freezing behavior in ChR2-injected animals during the initial two-minute baseline period across all three environments. Although there is no interaction, there was significant variance from the environment and the light-epoch (RM 2-way ANOVA with Tukey’s post-hoc correction; interaction: F_2.339,18.71_ = 1.977, p = 0.1615, Environment: F_1.952,15.61_ = 10.88, p = 0.0012, Light: F_1.172,9.38_ = 9.155, p = 0.0116, **Figure 1E**). The light-on epochs were further averaged and we saw the greatest effect on light-induced freezing in only the Small Box condition (RM one-way ANOVA with Tukey’s post-hoc correction; F_1_,_15_ = 8.239, p = 0.0039, **Figure 1E inset**). Importantly, this relationship between environment and light-induced freezing is not seen in control animals. We generated a difference score to compare overall light-induced freezing and found significant variation across virus conditions, but not the environment or an interaction between the two terms (RM two-way mixed-effects model with Tukey’s post-hoc correction; interaction: F_2,42_ = 2.111, p = 0.1338, virus: F_1,21_ = 7.320, p = 0.0132, environment: F_1.59,33.38_ = 2.938, p = 0.0775, **Figure 1F**).

Since the Large Box has dimensions commonly associated with Open Field assessments for anxiety, we examined if there was light-induced avoidance of the center in ChR2-injected mice. We found that there was a modest, albeit non-significant, decrease in the percentage of time spent in the center of the arena during light-on epochs in ChR2 animals (Sidak corrected t-test; t = 2.103, p = 0.0932; **Figure 1G, inset**). However, there was no significant variation in the amount of time spent in the center across virus conditions and light-epochs or an interaction between the two terms (RM two-way ANOVA with Sidak’s multiple comparisons for a main row effect; interaction: F_1,21_ = 1.991, p = 0.1729, virus condition: F_1,21_ = 0.5669, light epoch: F_1,21_ = 0.0755, **Figure 1G**). A difference score was also generated, and we saw no difference across groups (Welch’s unpaired t-test; t = 1.428, p = 0.1705, **Figure 1H**). Together, these data suggest that the same population of DG cells are not fixed to drive freezing per se but are capable of differentially driving behavioral responses in a manner contingent on the physical environment itself.

### Engram reactivation in the Small Box increases brain-wide cFos density

We next sought to measure brain-wide correlates of hippocampal CFC engram reactivation. Prior work suggests that artificially reactivating hippocampal engrams increases cFos expression in downstream areas that are implicated in mediating learning and affective states (Ramirez et al., 2015; Roy et al., 2022). We hypothesized that we would see significant differences in cFos expression in these regions as well as candidate regions that dictate behavioral outputs. In separate sets of animals, we tagged a hippocampal CFC engram as before but mice were subjected to only one day of reactivation in either the Small Box or the Large Box (**Figure 2A**). We sacrificed animals 90 minutes after the last light-on epoch to capture endogenous cFos expression and sent brain tissue for whole organ clearing, immunohistochemistry, light sheet microscopy, and cell quantification (LifeCanvas Technologies, Cambridge MA; see Methods). We first recapitulated light-induced freezing in ChR2 animals in the Small Box condition (RM two-way ANOVA with Sidak’s multiple comparisons for a main row effect; interaction: F_1,14_ = 10.62, p = 0.0057, virus condition: F_1,14_ = 3.785, p = 0.0721, light-epoch: F_1,14_ = 2.155, p = 0.1643, **Figure 2B + inset**), but saw no changes in the amount of time spent in the center in the Large Box condition (RM two-way ANOVA with Sidak’s multiple comparisons for a main row effect; interaction: F_1,14_ = 1.237, p = 0.2848, virus condition: F_1,14_ = 0.1337, p = 0.7201, light-epoch: F_1,14_ = 2.028, p = 0.1763, **Figure 2B + inset**). Over 100 brain regions were surveyed spanning anterior cortical areas to the posterior hindbrain (**Figure 2C, Table 1**). We first examined average cFos expression across 10 parent regions from the Allen Brain Atlas under each environmental condition. In the Small Box condition, where there was light-induced freezing, we observed that there were corresponding discoveries in the cortical plate (t = 4.147, p = 0.002), cortical subplate (t = 2.835, p = 0.0137) and the hypothalamus (t = 3.977, p = 0.002) in the ChR2 group (Multiple t-tests with 5% Benjamini-Hochberg FDR (Benjamini and Hochberg, 1995; Glickman et al., 2014); **Figure 2D**). Conversely, we found no discoveries in those parent regions under the Large Box condition (Multiple t-tests with 5% Benjamini-Hochberg FDR; **Figure 2E**). This supports the notion that optogenetic reactivation of a hippocampal engram is sufficient to drive cFos expression in a brain-wide pattern that itself depends on the environment in which stimulation occurred.

**Figure 2:**
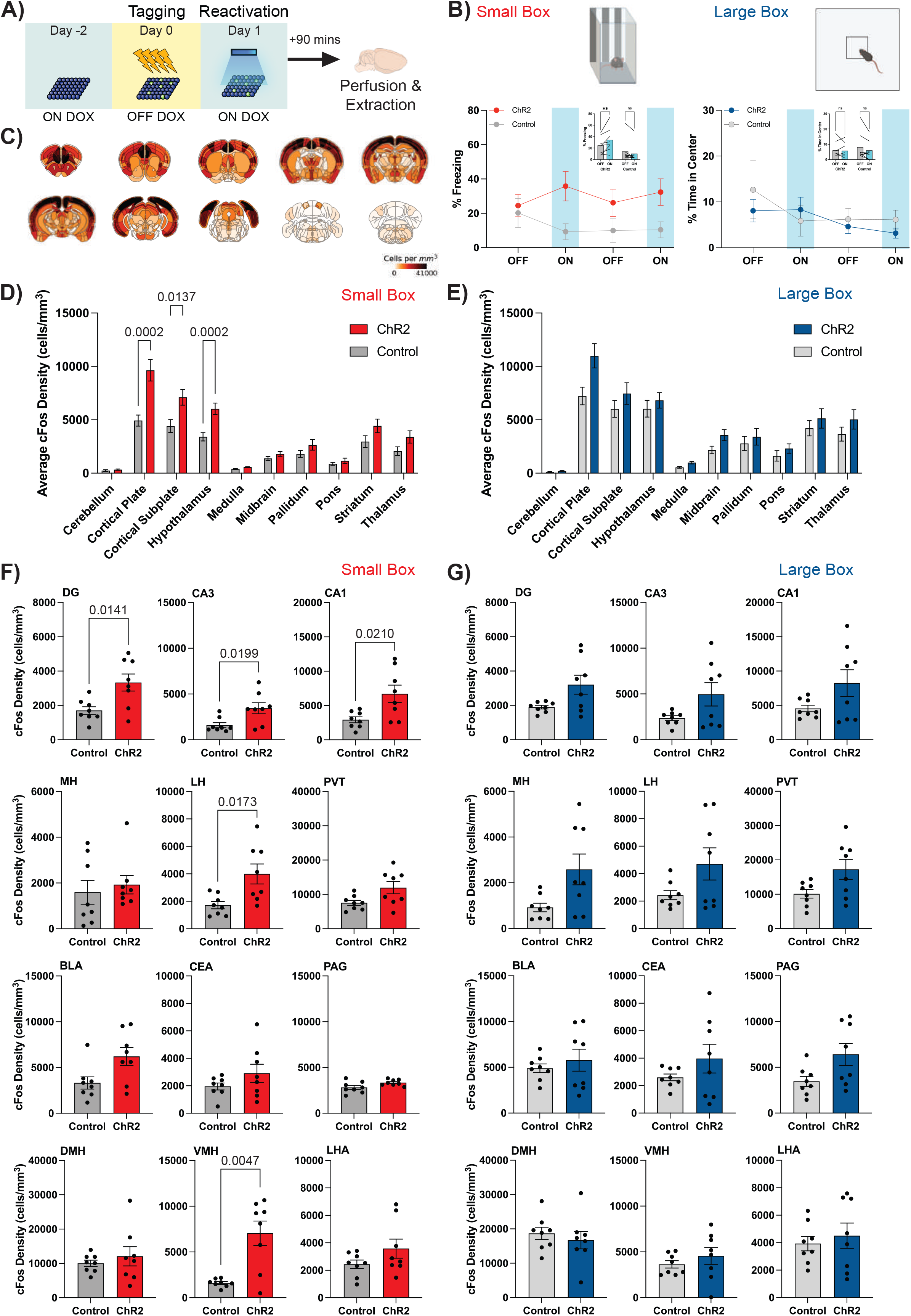
Engram reactivation in the Small Box increases brain-wide cFos density in areas mediating memory and behavior. (A) Schematic of experimental paradigm. Animals experience activity-dependent tagging of a CFC engram, but reactivation for only one day after the DOX window is closed. They are perfused 90-minutes after the last light-on epoch to capture peak endogenous cFos expression. (B) %Freezing levels for ChR2 animals (n = 8) and control animals (n = 8) for the Small Box condition across light epochs. There is a significant increase in the amount of light-induced freezing in only ChR2 animals (inset). %Time in Center for separate groups of ChR2 animals (n = 8) and control animals (n = 8) for the Large Box condition across light epochs. There is no difference in the amount of time in the center of the chamber across groups (inset). (**) p < 0.01. (C) Example heatmap of rodent brain-wide cFos density (cells/mm^3^). (D) Aggregation of the average cFos density in 10 parent brain areas registered to the Allen Brain Atlas in the Small Box condition. The observed p-values that are considered “discoveries” after the FDR correction are reported. (E) Aggregation of the average cFos density in 10 parent brain areas registered to the Allen Brain Atlas in the Large Box condition. The observed p-values that are considered “discoveries” after the FDR correction are reported. (F) cFos density for 12 individual regions of interest were compared between ChR2 and control animals in the Small Box condition using multiple unpaired Welch-corrected t-tests with a Benjamini-Hochberg correction of 5%. Observed p-values that are considered discoveries after the FDR are reported. (G) cFos density for 12 individual regions of interest were compared between ChR2 and control animals in the Large Box condition using multiple unpaired Welch-corrected t-tests corrected with a Benjamini-Hochberg correction of 5%. There were no observed p-values that were considered discoveries after the FDR correction. Data are presented as Mean ± SEM.

Next, we examined cFos density by individual brain regions. We honed in on twelve individual regions that have been heavily implicated in driving memory and defensive behaviors; specifically the hippocampus (DG, CA3, CA1; Fanselow and Dong, 2010; Scoviille and Milner, 1957), amygdala (BLA, CEA; Amano et al., 2011; Fadok et al., 2017; Herry and Johansen, 2014; Yu et al., 2016), the habenula (LH, MH; Pobbe and Zangrossi, 2010; Soria-Gómez et al., 2015; Stamatakis and Stuber, 2012; Zhang et al., 2016), the paraventricular nucleus of the thalamus (PVT; Do-Monte et al., 2015; Ma et al., 2021; Penzo et al., 2015), periaqueductal gray (PAG; Deng et al., 2016; Tovote et al., 2016), as well as the hypothalamus (DMH, VMH, LHA; Jardim and Guimarães, 2004; Jimenez et al., 2018; Kim et al., 2013; Lin et al., 2011). We predicted that these systems would be differentially engaged across conditions as a result of hippocampal CFC engram reactivation. In the Small Box condition, where we again observed light-induced freezing (**Figure 2B, left**), we found that there were discoveries in cFos density in the DG (t = 2.987, p = 0.014), CA3 (t = 2.779, p = 0.020), CA1 (t = 2.822, p = 0.021), LH (t = 2.926, p = 0.017), and the VMH (t = 4.043, p = 0.005) after applying a 5% FDR correction (**Figure 2F**). Interestingly, although there appeared to be enhanced cFos density in most of the selected regions in ChR2 animals, there were no discoveries in the Large Box condition when the FDR correction was applied (**Figure 2G**). This suggests that hippocampal CFC engram reactivation engages downstream areas that mediate memory and behavioral expression that is also contingent on the animal’s environment.

### Engram reactivation alters brain-wide cFos correlations

To understand the brain-wide interactions and functional relationships that occur during our engram manipulation, we constructed and analyzed cFos cell density networks using a custom pipeline freely available on GitHub (see Methods). We first visualized interregional Spearman’s correlations for all groups in all conditions and organized them based on the parent region in the Allen Brain Atlas. We observed significantly greater coactivation in the ChR2 conditions than in the control conditions (**Figure 3A-B**). Next, we used uniform manifold approximation and projection (UMAP; McInnes et al., 2020), a non-linear dimensionality reduction technique to represent the data in two-dimensional space. UMAP allows us to test whether our correlation values will segregate based on parent brain region or by group condition. Interestingly, when applying UMAP, we found that our data grouped by condition, such that the majority of the correlation values aggregated in a single cluster, and the values by group, irrespective of parent brain region (**Figure 3C-D).** Notably, while the control conditions were the closest in terms of their centers of mass, the ChR2 conditions clustered away from each other. This lends credence to the idea that our environmental and engram manipulations create distinct brain states.

**Figure 3:**
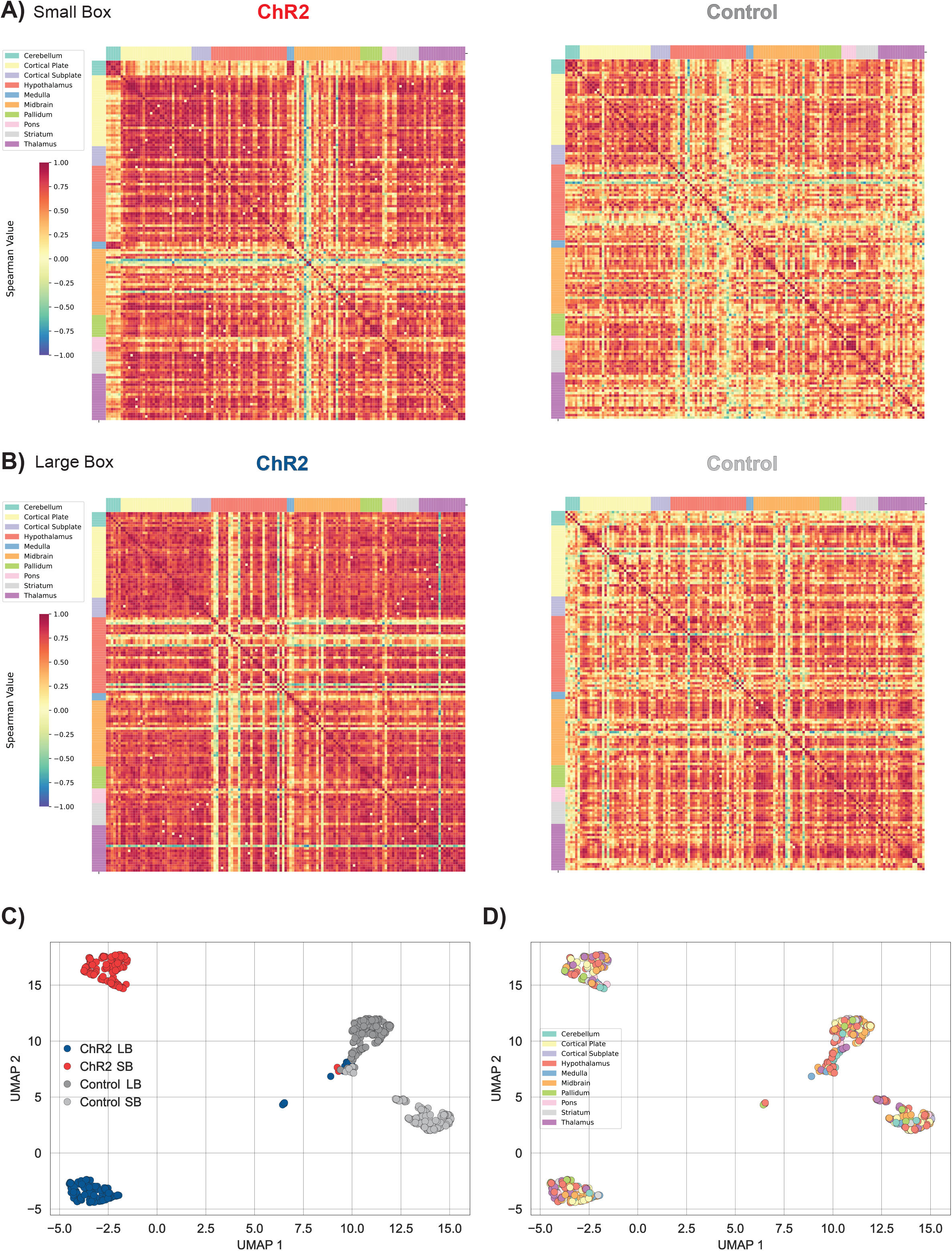
Engram reactivation differentially increases cFos correlations across environments. (A) cFos correlation matrices organized by Allen Brain Atlas anatomy generated from the Small Box condition. There are greater positive Spearman correlations in the ChR2 group (red) when compared to Control. (B) cFos correlation matrices organized by Allen Brain Atlas anatomy generated from the Large Box condition. There are greater positive Spearman correlations in the ChR2 group (red) when compared to Control. (C) UMAP representations of ChR2 and Control groups across environmental conditions (i.e., Small Box [SB]; Large Box [LB]). All four conditions were separated in linear space, yet both Control groups were closer in space than either of the ChR2 groups, showing that these states are inherently distinct. (D) UMAP representations of 147 brain regions spanning Allen Brain Atlas parent regions (legend). Brain regions do not show distinct segregation, as all of these colors are intermingled, suggesting that engram stimulation does not bias particular brain regions into separable linear spaces.

### Engram reactivation induces distinct topological features in brain-wide networks

To understand how our engram manipulation induced changes to the functional pairwise relationships between brain regions, we modeled our correlation data using graph theory. We defined graphs such that the aforementioned correlation data would serve as our adjacency matrices, except we thresholded the Spearman coefficients such that only the top 25% were kept (**Figure 3A-B**). This ensured that when we calculated network statistics, differences were not due to edge densities across networks (Garrison et al., 2015; Rubinov and Sporns, 2011). Our graphs were then defined such that the brain regions formed the set of nodes and the Spearman correlation coefficients were the set containing the edges between nodes. We first asked how different regions of the brain are compartmentalized together across our control and experimental groups. To this end, we used Louvain community detection (Blondel et al., 2008) to functionally segregate our networks into local communities–that is, groups of nodes that are highly connected within a community, and less densely connected between communities. Interestingly we noticed that each of our four conditions formed primarily four large communities, and a varying amount of significantly smaller communities, with the largest nonprimary partition consisting of only three nodes (**Figure 4A-B)**. Next, in the ChR2 Small Box network, we saw that the largest community partitioned regions highly associated with memory and fear responses, such as the DG, CA1, CA3, thalamic, and hypothalamic subregions, and this was the only condition in which all components of the trisynaptic circuit are within one community (**Figure 4A**). Our data here may indicate that while the community structures are similar in nature across conditions, engram reactivation may reorganize communities in a way that could mediate memory-guided behavioral responses.

**Figure 4:**
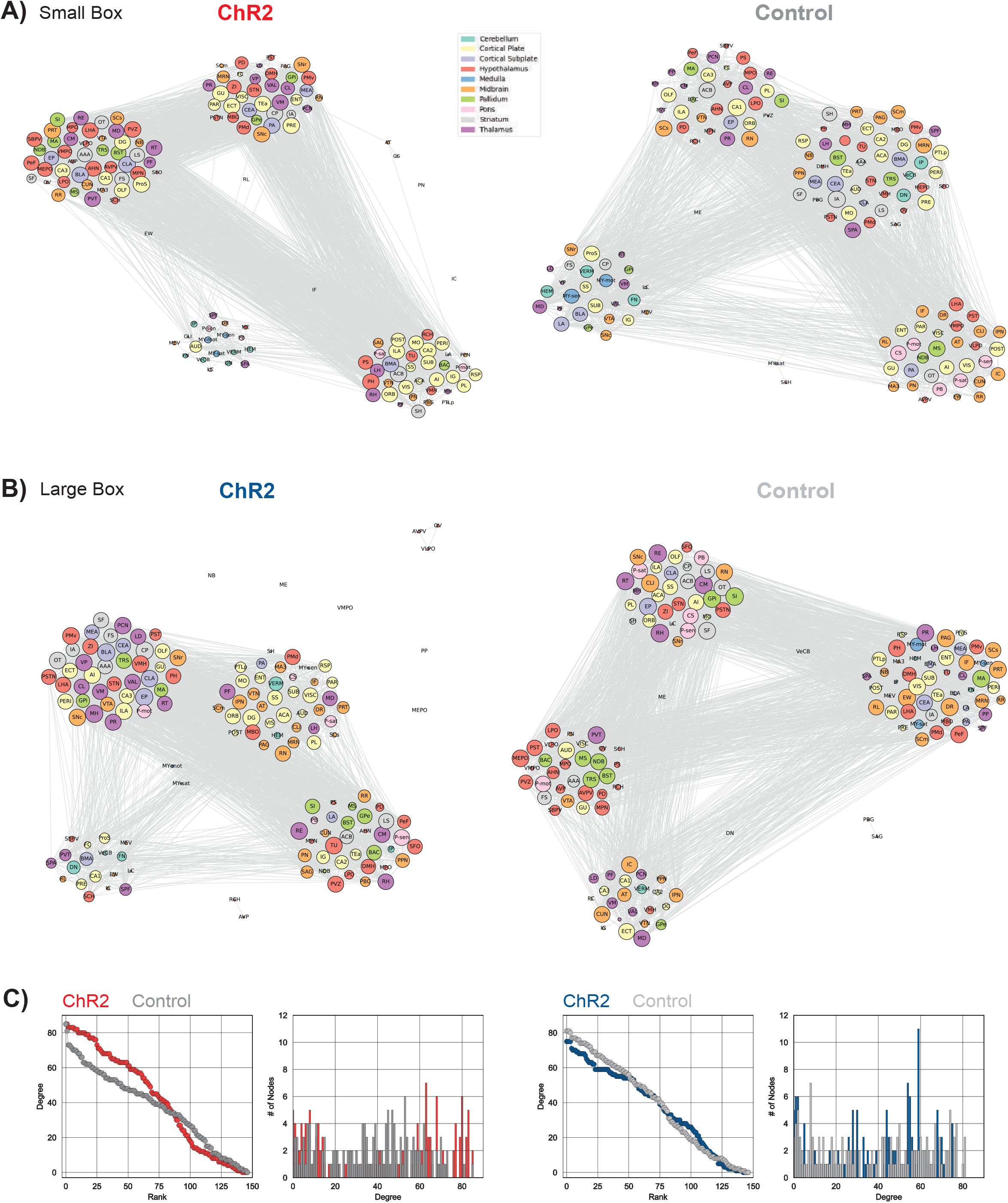
Engram reactivation creates unique network topologies. (A) ChR2 (Left) and Control (Right) networks generated in the Small Box condition after edge thresholding was applied. Both networks have four main communities, yet the composition of Allen Brain Atlas regions in each community is different. (B) ChR2 (Left) and Control (Right) networks generated in the Large Box condition after edge thresholding was applied. Both networks have four main communities yet the composition of Allen Brain Atlas regions in each community is different. Additionally, there are disconnected components in the ChR2 condition as a result of edge thresholding. (C) Degree-rank distribution across all 147 brain regions for all conditions. There are more nodes of lower degree rank in the ChR2 Small Box condition when compared to Control. Yet, there are no differences in degree rank under the Large Box condition.

We next plotted the degree-rank plots for each condition to measure how similar the control and experimental conditions were in terms of their degree distributions, and found that the Large Box groups were more similar than the Small Box groups, which had greater separation between the two curves (**Figure 4C)**, further evidencing differences in network topology. From there, we calculated common centrality metrics: degree centrality (i.e., number of edges connected to a node), betweenness (i.e., how often a node is in the shortest path between all pairs of nodes), closeness (i.e., average length of the shortest possible path between a node and every other node in the graph), clustering coefficient (i.e., how often a node is connected to another node that forms a local clique), and eigenvector centrality (i.e., how often a node is connected to highly connected nodes) (**Figure 5A, top)**. We first observed that there were no differences between the betweenness and eigenvector centrality of any of the conditions, indicating that the average node in either network is not more connected to highly connected nodes, nor is it on the shortest path between any other two nodes (**Figure 5A, top)**. However, in both closeness and degree centrality, the ChR2 Small Box and Control Large Box conditions were both significantly greater than the Control Small Box condition, indicating that on average the nodes in these networks were more connected and had a shorter path length to every other node (**Figure 5A, top)**. The largest differences, however, were seen when comparing the clustering coefficients: the ChR2 Small Box condition had a significantly higher average coefficient than all other conditions, suggesting that these nodes were more likely to be connected to nodes that themselves were near or completely connected to their adjacent nodes (**Figure 5A, top)**. Additionally, the Control Large Box had a statistically higher clustering coefficient than the Control Small Box condition. From these data we suggest that engram reactivation induces alterations in network structure that is unique across conditions.

**Figure 5:**
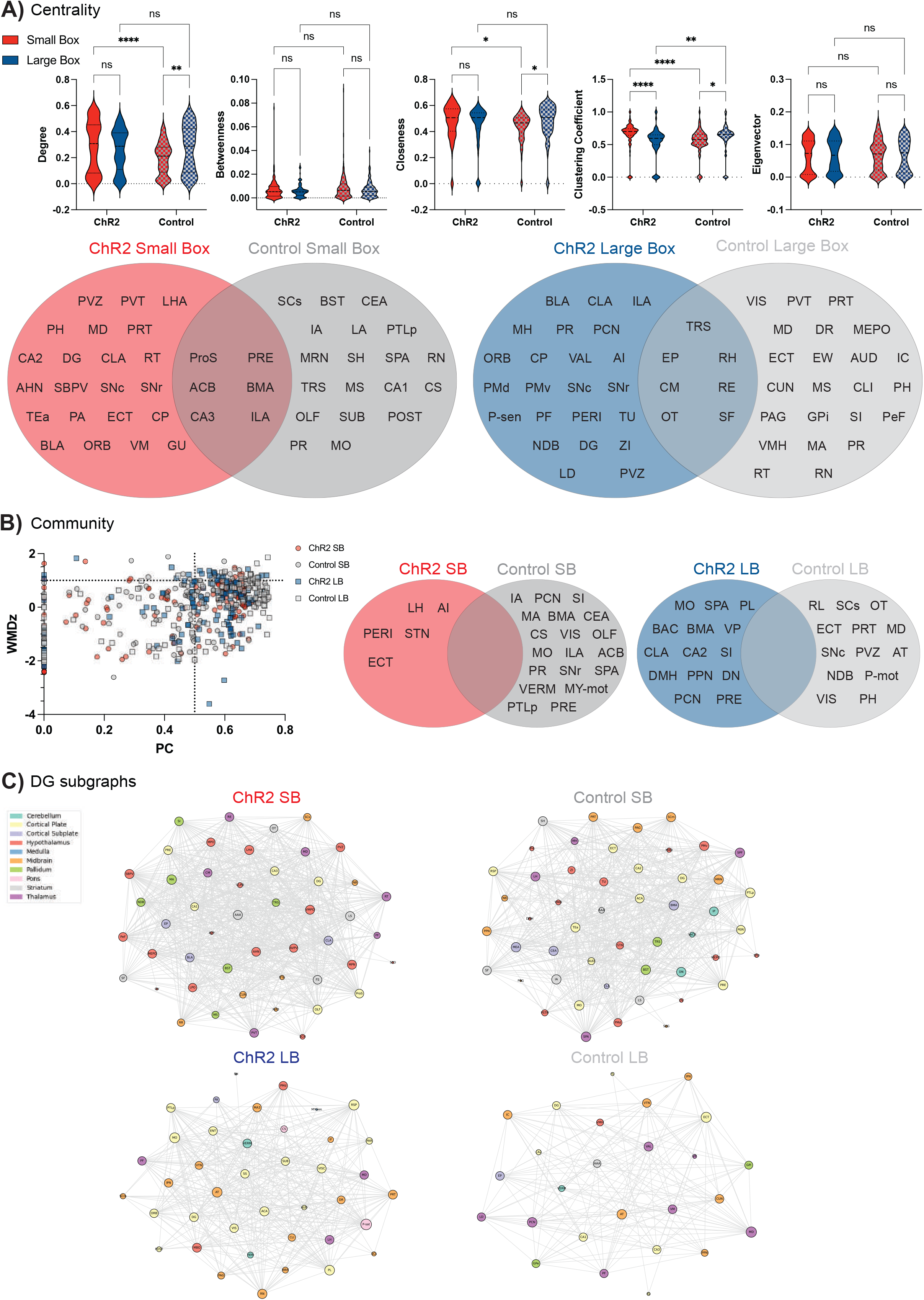
Regions involving memory and behavior act as putative hubs in ChR2 networks. (A) Degree, Betweenness, Closeness, Clustering Coefficient, and Eigenvector centrality metrics are compared across all four experimental conditions after network edges were thresholded by the strongest 25% of edges (Top panel). (*) p < 0.05, (**) p < 0.005, (****) p < 0.0001. “Hub scores” were generated for all 147 nodes in the network. Nodes falling into the top 20% (Degree, Betweenness, Closeness, Eigenvector) or bottom 20% (Clustering Coefficient) received a +1, and central hubs were identified as having a score of 3 or above (Bottom panel). (B) Within-module degree z-score (WMDz) and participation coefficient (PC) were generated for all 147 regions under all four experimental conditions. Classification of a community-based hub would pass a WMDz threshold of 1.0 and a PC threshold of 0.5 and nodes were identified as such for each condition. (C) DG-containing subgraphs were generated under each condition to identify nodes that are clustered in the same module after the Louvain clustering algorithm was applied.

### Engram reactivation differentially engages hubs across environmental conditions

Hubs in a network are nodes that are of high importance for signal propagation and are defined using a series of centrality metrics in addition to community-based characteristics when clustering algorithms are applied (Guimerà and Nunes Amaral, 2005; van den Heuvel and Sporns, 2013). To identify nodes that act as central hubs, we generated a “hub score” by characterizing the node distribution across five centrality metrics: degree centrality, betweenness, closeness, eigenvector centrality, and clustering coefficient (Coelho et al., 2018; van den Heuvel et al., 2010). Nodes that fell in the top 20% (or bottom 20% in the case of clustering coefficient) of any of these metrics receive a +1 added to their hub score. Here, we defined central hubs as having a score of 3 or greater (**Figure 5A, bottom**). We found 22 central hubs unique to the ChR2 Small Box network, with many implicated in memory (e.g., DG, ECT, CA2, BLA) and behavior (e.g., PVT, LHA; see **Figure 5A, bottom**). Interestingly, all six of the shared hubs between the ChR2 and Control Small Box networks are important for memory-guided actions. We speculate that these hubs may be involved in processing shared representational features of the Small Box environment through their coordinated activity. There were 23 central hubs in the ChR2 network generated under the Large Box condition, many of which are unique to the network. Importantly, the DG and BLA are two shared hub regions in both ChR2 networks generated from the Small Box and Large Box condition, demonstrating that our engram manipulation does engage largely characterized memory systems. Collectively, these findings provide an extensive list of candidate hubs that are differentially recruited across environments, furthering the evidence that these networks are distinct.

Application of clustering algorithms functionally segregates nodes based on shared correlation composition across the network. Thus, we further characterized nodes by how they communicated within and across communities. For each node, we generated a respective within-module degree z-score (WMDz) and participation coefficient (PC; (Guimerà and Nunes Amaral, 2005). We defined community-based hubs as having both a WMDz score of 1.0 or above and PC score of 0.5 or higher (**Figure 5B**). There were only four unique communitybased hubs in the ChR2 Small Box network. Surprisingly, we found that there are no shared community-based hubs across ChR2 and control networks in either environment, further suggesting that the community organization is unique to each condition. Furthermore, we then determined the differences in the DG-containing community (i.e., DG subgraph). As our optogenetic approach targeted engram cells contained within the DG, the respective subgraph containing connected edges could elucidate what changes our perturbation directly causes to neighboring nodes (**Figure 5C**). We found more hypothalamic, thalamic, and pallidum areas in the DG subgraph in the ChR2 Small Box network. Conversely, areas spanning the cortical plate predominated the ChR2 Large Box DG-containing subgraph. Overall, these results qualitatively show that there are unique patterns of co-activation as a result of DG-centered CFC engram reactivation.

### Chemogenetic inactivation of the lateral hypothalamus during engram reactivation alters behavior

Our brain-wide network data pointed to the lateral hypothalamic area (LHA) as a central hub in the ChR2 networks from the Small Box condition. The LHA was also close to meeting criteria for acting as a community-based hub (WMDz = 0.95, PC = 0.62), and was central to the DG-containing subgraph of the ChR2 Small Box condition. In support, recent studies suggest that circuits involving the LHA are implicated in mediating fear and anxiety-like behaviors (Bonnavion et al., 2015; Concetti et al., 2020; Jimenez et al., 2018; Kim et al., 2013); thus, we tested whether or not the LHA was necessary for light-induced freezing.

A new set of experimental mice containing activity-dependent ChR2 in the DG also co-expressed a pan-neuronal inhibitory hM4Di in the LHA. Control animals contained activity-dependent ChR2 in the DG but an empty vector in the LHA (i.e., mCherry only). All animals were subjected to CFC during the tagging phase as described above. The next day, all subjects received an intraperitoneal injection of clozapine-n-oxide (CNO; 3 mg/kg) 30-minutes prior to hippocampal CFC engram reactivation in the Small Box (**Figure 6A**). Mice in the hM4Di group exhibited near floor-levels of freezing when compared with control animals (RM two-way ANOVA with Sidak’s multiple comparisons for a main row effect; interaction: F_1,10_ = 0.1570, p = 0.7002, virus condition: F_1,10_ = 9.387, p = 0.0120, light epoch: F_1,10_ = 0.8797, p = 0.3704, **Figure 6B + inset**). Although the control animals had ChR2 in tagged DG cells, we did not observe any light-induced freezing across epochs (t = 0.9434, p = 0.6002). We speculate that the lack of light-dependent freezing may be attributed to off-target effects due to clozapine metabolism or due to an occlusion effect, as freezing was sustained at ~30% for the test in control animals. Therefore, to gain histological insight, we next measured the effects of CNO administration of endogenous cFos expression in the LHA by the number of cFos cells after CFC engram reactivation. Surprisingly, we saw a trending, although non-significant, increase in cFos cells in the LHA in hM4Di-expressing animals when compared to controls (Welch’s unpaired t-test; t = 2.165, p = 0.0558; **Figure 7C**), which underscores the complex relationship between chemogenetic inhibition and the ensuing local cellular activity. Although our experiments here target the LHA, other previously identified hubs are also suitable candidates for future interventional studies.

**Figure 6:**
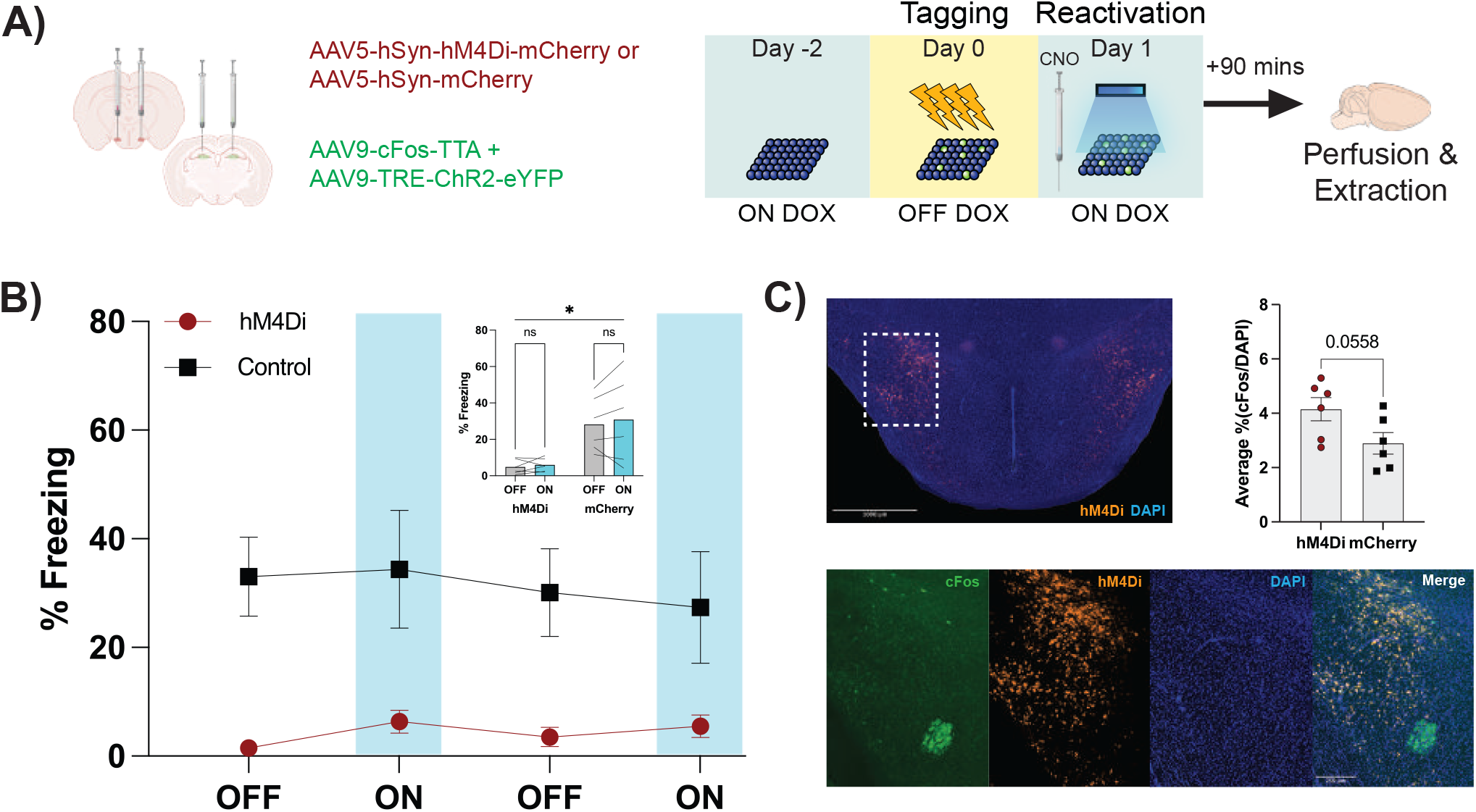
Chemogenetic inactivation of the LHA produces alterations in behavior but not cFos expression. (A) Experimental design for inactivation of the LHA during CFC engram reactivation. Animals underwent surgical procedures for two viral infusions of activity-dependent tagging of engram cells and hM4Di administration in the DG and LHA, respectively. The behavioral design follows those outlined in Figure 2 but with CNO administration 30 minutes prior to CFC engram reactivation in only the Small Box condition. (B) %Freezing levels for all animals (hM4Di, n = 6; Control, n = 6) across light epochs. There is a significant effect of virus condition in the amount of freezing throughout the experimental session, as hM4Di animals exhibited near-floor levels of freezing behavior. Yet there are no significant differences across light epochs within each group (inset). (*) p < 0.05. (C) Histological representation of LHA targeting (Top left panel; scale bar represents 1000 μm), virus expression, and cFos (Bottom panel; scale bar represents 250 μm). Cell quantification revealed greater, but not significant, endogenous cFos expression in the hM4Di-injected animals than controls (Top right panel). All data presented as Mean ± SEM.

**Figure 7:**
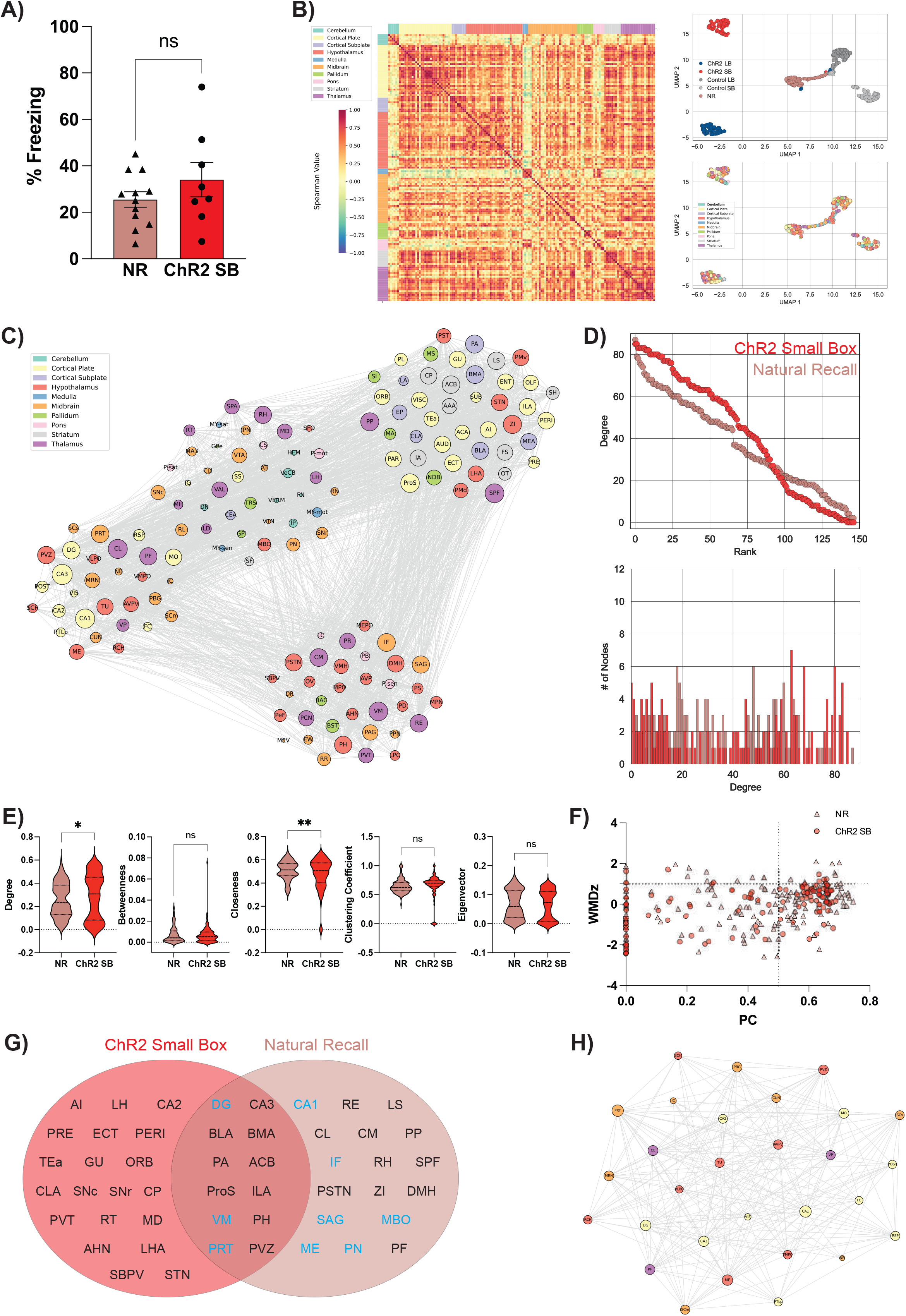
Natural and artificial memory recall are distinct in network features, yet share similar behavioral output and hubs. (A) %Freezing levels for both Natural Recall (“NR”; n = 12) and the ChR2 Small Box condition (“ChR2 SB”; n = 8; Average Light-on). There was no statistically significant difference in freezing behavior between the two groups. (B) cFos correlation matrix generated from the Natural Recall group (Left). UMAP representations of Natural Recall and other groups as represented in Figures 3C-D (Top right). Natural Recall correlations clustered into a distinct linear space in between ChR2 and Control groups, but were slightly closer to Control groups. Allen Brain Areas were also not segregated (Bottom right). (C) Network representation of Natural Recall. There are four communities with unique Allen Brain Area compositions. (D) Degree rank plots show that there are more lower ranking nodes in the ChR2 Small Box group and higher ranking nodes in the Natural Recall group. (E) Violin plots comparing five centrality metrics across groups. There is a significant difference in the average degree centrality and closeness. (F) Within-module degree z-score (WMDz) and participation coefficient (PC) were generated for all 147 regions for Natural Recall and ChR2 Small Box (SB). Classification of a community-based hub would pass a WMDz threshold of 1.0 and a PC threshold of 0.5 and nodes were identified as such for each condition. (G) Both central and community-based hubs are listed between both conditions. There are shared hubs spanning regions highly implicated in mediating memory-guided actions and hubs distinct to either ChR2 Small Box or Natural Recall. Community-based hubs for the Natural Recall condition from F are highlighted in blue. (H) DG-containing subgraph from the Natural Recall condition. All subregions of the hippocampus and regions spanning thalamic, hypothalamic and midbrain pathways are housed in the community containing the DG.

### Natural recall and engram reactivation in the Small Box exhibit shared features

Previous works using network analyses have examined cFos correlations in areas spanning putative memory and defensive behavior pathways after fear memory recall (Cho et al., 2017; Vetere et al., 2017; Wheeler et al., 2013). Thus, we predicted that our network generated by engram reactivation in the Small Box condition would demonstrate shared features with the natural recall of a fear memory. To that end, we compared the networks resulting from optogenetic engram reactivation to natural conditions in which animals recalled a bona fide fear memory. Naive mice were subjected to CFC as previously described, and placed back into the original context a day later before examining brain-wide cFos correlations. Behaviorally, the Natural Recall group and the ChR2 Small Box group froze at statistically comparable levels (Unpaired t-test; t = 1.184, p = 0.2519; **Figure 7A).** Interestingly, despite displaying similar freezing levels, we found a mix of shared and differing network characteristics.

We first observed that, overall, the Spearman correlation matrix for the Natural Recall group contained fewer high magnitude positive Spearman coefficients. Furthermore, when plotting into UMAP space, the Natural Recall group was located between both optogenetic and control states, possibly indicating that Natural Recall had shared characteristics between the two conditions. When comparing the network statistics, only degree and closeness were significantly different: the ChR2 Small Box had higher degree nodes, but Natural Recall had nodes with higher closeness. Thus, the group receiving optogenetic stimulation in the Small Box had more connections, but the Natural Recall condition had nodes that were closer spatially **(Figure 7E),** indicating that the ChR2 Small Box network was more connected but the individual nodes of the Natural Recall group were more similar. Additionally, the degree-rank plot showed that the ChR2 Small Box group had more nodes with greater average degree, indicating a more globally connected network (**Figure 7D).** Since the clustering coefficient was larger in the ChR2 Small Box group, but not in the other networks, and was not significantly different from Natural Recall, this could indicate that the average clustering coefficient of a network we speculate can be used to predict fear states in this context **(Figures 5A, 7E).** We also compared the hubs between these groups and found they shared more regions than any other pair of conditions **(Figure 7F-G).** As predicted, these groups shared the DG, CA3, BLA, and other regions which have been extensively implicated in memory and fear responses (Fendt and Fanselow, 1999; Gross and Canteras, 2012; Silva et al., 2016; **Figure 7G).** Notably, these are the only two networks which contained the trisynaptic circuitry (DG, CA1, CA3) within the same community detected through Louvain community detection (**Figure 7C),** while in the artificial recall condition, more hypothalamic areas were in the same community as these hippocampal regions **(Figures 5C, 7H).** Overall, these results imply that shared behavioral phenotypes do not necessitate identical brain states, indicating that memory generated fear states can have differential patterns of brain activity.

## DISCUSSION

These results demonstrate that hippocampal CFC engrams have the ability to drive behaviors in a manner dependent on the physical environment. Furthermore, artificial reactivation of a hippocampal CFC engram across different environments induced whole-brain activity in a manner topologically distinct between environments, and network hub regions mediating memory and behavior play a more active role in our experimental groups.

Our behavioral findings suggest that light-induced freezing becomes more apparent as the size of an environment becomes constrained. Previous studies examining innate fear responses during TMT exposure also show this relationship between the environment size and freezing (Rosen et al., 2008; Wallace and Rosen, 2000). Rodents exposed to TMT in smaller arenas often defaulted to freezing; others opted for more active ambulatory responses such as avoidance as environments were less spatially constrained. Together, these data demonstrate that both external (e.g., TMT exposure) and internal stimuli (e.g., HPC engrams) engage in differing behavioral outputs depending on the physical layout of the environment. In support, recent research suggests that activated fear engrams are sufficient to drive active avoidance-like behaviors (Chen et al., 2019; Ramirez et al., 2013; Redondo et al., 2014). Unlike previous work, which uses separate cohorts of animals for separate environmental conditions, our results demonstrate within-animal modulation of behavior during engram reactivation under different environmental size contingencies. Specifically, our data shows that while these same cells drive freezing in the Small Box condition, the lack of freezing observed in the Large Box condition underscores the behavioral flexibility driven by activated DG cells.

Future studies can examine the effects of hippocampal CFC engram reactivation and light-induced freezing in more complex environmental conditions such as introducing an exit or shelter that warrants the rodent to engage in more navigation-based escape strategies (Mangieri et al., 2019; Salay et al., 2018; Vale et al., 2017; Wang et al., 2021). We predict that the latency for seeking either an exit or shelter would decrease at the onset of hippocampal CFC engram reactivation. Additionally, while our study measured freezing as a proxy of artificial fear memory recall, a negative affective state such as fear can manifest in a variety of behaviors and across sexes (Blanchard and Blanchard, 1989; Fanselow et al., 1987; Colom-Lapetina et al., 2019). There are many defensive strategies that encapsulate the canonical “fight, flight, or freeze” behaviors which animals implement depending on their situation (Bolles 1970; Blanchard and Blanchard, 2008; Grossen and Kelley, 1972). These behaviors also include tail-rattling as a sign of defensive aggression (Salay et al., 2018), risk assessment via stretch-attend posture (Molewijk, 1995), jumping as a means of escape (Mangieri et al., 2019), and selfgrooming as a de-arousal technique after an aversive experience (Kalueff et al., 2016; Song et al., 2016). Additionally, certain defensive behaviors such as darting are sex-dependent in dimorphic species such as rodents (Gruene et al., 2015; Shansky and Murphy, 2021). Thus, in addition to freezing, we speculate that animals may engage in these behavioral modalities during hippocampal CFC engram reactivation under different environmental conditions which may also be sex-dependent in nature. Future studies can examine the behavioral effects of hippocampal CFC engram reactivation in females across environments to test for biological sex mediating behavioral strategies; we predict that female mice will exhibit a mix of light-induced freezing and darting responses in the Small Box, but light-induced darting will predominate in the Large Box. Indeed, there are a variety of interleaving brain-wide pathways that mediate alterations in behavioral strategies (Fanselow, 1994; Fendt and Fanselow, 1999; Gross and Canteras, 2012; Silva et al., 2016; Tovote et al., 2015), which points to the capacity for hippocampal CFC engrams to produce changes in internal brain states.

Our brain-wide analyses revealed that optical stimulation of a hippocampal CFC engram is capable of globally increasing cFos expression. This dovetails with recent work in the field showing that 4 Hz stimulation of a CA1 engram ensemble recruits regions downstream in a brain-wide manner (Roy et al., 2022). Furthermore, these changes coincide with greater positive Spearman correlations, indicating that our optical stimulation is coordinating the brain’s activity at the network level. This suggests that artificial memory activation pushes global brain activity to a state of greater functional connectivity. Interestingly, however, the levels of freezing between natural and artificial recall in the Small Box condition were not significantly different. We believe that the brain can be in two different physiological states that may not be fully captured by freezing alone. This is further supplanted by our UMAP results, which showed that brain regions segregated not by their parental region (e.g., cortical subplate, hypothalamus, etc.), but rather by their experimental condition, suggesting that each one of our paradigms elicited a unique pattern of brain activity.

Through our community detection analyses we probed which regions of the brain were more densely connected to one another. One of the most interesting results of this analysis was that the trisynaptic circuitry was within the same partition for the Natural Recall and ChR2 Small Box groups. This could explain why these two groups froze at statistically significant levels compared to the other conditions, as the hippocampus was functionally recruited similarly for both, perhaps similar states of global coactivity to drive freezing responses. Of note, many hypothalamic, thalamic, and pallidum regions were in the ChR2 Small Box DG subgraph, yet the one from the Natural Recall group contained mostly cortical plate regions with some hypothalamic, thalamic, and midbrain regions, which could point to our optogenetic paradigm inducing a heightened state of fear, even without manifesting as a further increase in freezing. As all of the conditions created primarily four main communities with smaller partitions of varying size, this could point to alterations in the relevant activity patterns across the brain becoming coactive in a task-specific manner.

In graph theoretical analyses, hubs are defined as nodes that are crucial to mediating signal propagation throughout the network. We found more than 20 hubs under each experimental condition, some of which were shared, but many which were unique. This indicates that there are many brain-wide regions that are important for maintaining the structure of our generated networks under different conditions such as environment size. Interestingly, we found that the hub composition for the ChR2 Small Box condition recruited many areas that support learning (e.g., DG, CA3, BLA) and behavior (e.g., ACB, PVT, LHA). Additionally, many of these hubs are also shared with ones found in the Natural Recall network (e.g., DG, CA3, BLA, ACB). Based on this shared hub composition, we posit that our engram manipulation stimulates the brain into a memory-driven fear state, similar to natural memory recall, even if other network features are partly distinct across conditions. Although our *in vivo* hub inhibition in the ChR2 Small Box focused on the LHA, as it was a unique hub to the network, future work may determine the relative importance of each of these 20+ hubs for fear-induced freezing in both artificial and natural recall (Chen et al., 2019; Vetere et al., 2017). Taken together, our work bridges both biological- and network-based approaches for studying memory and behavior.

## METHODS

### Animals

Wildtype males (2-3 months of age; Charles River Laboratories) were housed in groups of 3-4 per cage. Aggressor mice were separated from cagemates as needed and were single housed with extra enrichment. All mice were kept on 12:12 light-dark cycles (0700-1900) in humidity-controlled colony rooms and had *ad libitum* access to standard rodent chow and water. Upon arrival at the facility, mice were left undisturbed for three days. We substituted the rodent chow with 40 mg/kg doxycycline (DOX) chow 24-hours prior to surgery. Surgerized mice were left undisturbed for ten days to recover. All subjects were treated in accordance with protocol 201800579 approved by Boston University’s Institutional Animal Care and Use Committee (IACUC Protocol 17-008).

### Viruses

The two viruses for the activity-dependent tagging system were packaged at the University of Massachusetts Amherst Viral Vector Core. The first virus contains a pAAV-cFos-tTA plasmid vector and the second a pAAV-TRE-ChR2-eYFP (or TRE-eYFP for control experiments). Neural activity induces the expression of the AAV9-cFos-tTA construct and generates tTA proteins in cells. The tetracycline transactivator (tTA) proteins then bind to the tetracycline response element (TRE) to induce the expression of either -ChR2-eYFP or -eYFP. This system is regulated by doxycycline (DOX), a tetracycline derivative, for strict temporal control over when cells are labeled. The removal of DOX from the system opens a tagging window to allow tTA to bind to TRE. For chemogenetic inactivation experiments, we used either an AAV5-hSyn-hM4D(Gi)-mCherry (Addgene; 50475-AAV5) or AAV5-hSyn-mCherry (Addgene; 114472-AAV5).

### Surgical Procedures

Mice were placed in an induction chamber and anesthetized with a mixture of 4% isoflurane and 70% oxygen and were maintained at 2% isoflurane when mounted in the nosecone of the stereotaxic frame (Kopf Instruments, Tujunga, CA, USA). We applied ophthalmic ointment to both eyeballs to prevent them from drying during surgery. Hair was removed via topical application of hair removal cream, and the scalp was cleaned with ethanol and betadine. We then applied 2% lidocaine (Clipper Distributing Company) to the surface of the scalp for topical analgesia. We made a scalp incision to expose the skull. Peroxide was applied to the surface of the skull to bleach the skull sutures and the skull was then leveled between bregma and lambda. Bilateral craniotomies were made above the site of viral injection.

All injection coordinates are in relation to bregma (in mm): for the dentate gyrus (DG), AP = −2.2, ML = ±1.3, DV = −2.0; for the lateral hypothalamic area (LHA), AP = −1.3, ML = ±1.25, DV = −5.3. A 33-gauge beveled needle connected to a 10 μl Hamilton syringe attached to a micro-infusion pump (UMP3, World Precision Instruments) was used for the viral injections. The needle was lowered 0.2 mm past the injection site and was kept stationary for 2 minutes. We then raised the needle to the site of injection and waited one minute before virus infusion. We bilaterally injected the DG with 200 nl of the AAV9-cFos-tTA & AAV9-TRE-ChR2-Venus viral cocktail at a rate of 110 nl/min. Five minutes after the injection was complete, we then moved the needle 0.2 mm above the injection site and waited another three minutes before complete needle removal. DG-injected mice were then implanted with bilateral fiber optics (200 μm core diameter; Doric Lenses) directly above the site of viral injection (−1.8 DV). In separate cohorts of animals undergoing DG + LHA surgery, we additionally injected 300 nl of AAV5-hSyn-hM4D(Gi)-mCherry or AAV5-hSyn-mCherry bilaterally at a rate of 110 nl/min. Skull screws were anchored for implant support. We applied layers of metabond and dental cement (A-M Systems) to create a cap over the skull. All mice received a 0.1 ml intraperitoneal injection of 0.3 mg/ml buprenorphine, a subcutaneous 0.5 mg/ml injection of meloxicam, and 0.2 ml of subcutaneous saline following surgery and were placed on a heating pad until conscious recovery.

### Fear Conditioning and Tagging

All mice were handled for two days prior to experiments. After the second day of handling, the doxycycline (DOX) diet was swapped with rodent chow and the animals were left undisturbed for two days prior to contextual fear conditioning (CFC). This opened the window for activity-dependent viral labeling during the CFC tagging period. All mice were placed in conditioning chambers with plexiglass walls and a grid floor (Coulbourn Instruments, Whitehall, PA, USA). This grid floor was connected to a precision-shocker and delivered a series of four foot shocks (2 sec, 0.5 mA intensity) throughout the duration of the 8-minute tagging session. Mice were placed back on the DOX diet in a clean cage immediately after tagging and remained on DOX for the duration of the experiment. Video data was collected via overhead cameras (Computar) that interface with FreezeFrame (Actimetrics, Wilmette, IL, USA). FreezeFrame can both control the delivery of the foot shocks and perform rudimentary freezing analyses. Freezing during CFC was defined as bouts of 1.25 secs or longer with minimal changes in pixel luminance as defined by a numeric pixel threshold, N.

### Optogenetics

We tested each patch cord before optogenetic experiments to ensure that each patch cord generated at least ~10 mW of power. Fiber optic implants were plugged into a patch cord connected to a 450 nm laser diode (Doric Lenses). For the large, medium, and small arena sessions, mice were allowed to freely explore for ten minutes. Each session began with a two-minute baseline period followed by two duty cycles of optogenetic stimulation. Each duty cycle began with a two-minute light stimulation (light-on; 20Hz, 10ms pulses) followed by two minutes of no light (light-off). At the end of the final light-off epoch, mice were unplugged from the patch cord and returned to their home cage.

### Chemogenetics

For chemogenetic silencing of the lateral hypothalamus, we used the inhibitory Designer Receptor Exclusively Activated by Designer Drugs (DREADDs), hM4Di, fused to the human synapsin (hSyn) reporter. The hM4Di drives inhibition of infected neurons when bound to clozapine-n-oxide (CNO; SigmaAldrich). A 0.6 mg/ml solution containing CNO was prepared in sterile saline and 0.5% dimethyl sulfoxide. All mice were intraperitoneally injected with saline for five days prior to the OFF DOX period for acclimatization. On the day of chemogenetic experiments, animals were then injected with a 3 mg/kg dose of the CNO solution 30 minutes prior to the optogenetic reactivation of the CFC engram.

### Behavioral Assays

All behavioral assays were conducted during the light cycle (0700-1900). During this time, mice had *ad libitum* access to DOX chow (or regular chow for the Natural Recall group) and water. Any noticeable aggressor mouse was separated to prevent any injury to cagemates.

#### Large Box

The large environment was a 63 cm L x 63 cm D x 45.5 cm H arena with opaque walls and a white matte bottom. The center of the chamber was demarcated with a 32 cm x 32 cm square. We additionally introduced orange scent and dimmed the overhead lighting as new contextual information. Mice were allowed to freely explore for 10 minutes. Optogenetic stimulation was delivered as described above.

#### Mid Box

This experiment was conducted in a conditioning chamber normally suited for rats (12 in L x 10 in D x 12 in H; Coulbourn Instruments). We taped laminated sheets of paper with a cross-hatched design to the walls and placed laminated paper with a vertical bar pattern on the floor to eliminate any contextual similarities. We additionally introduced a vanilla scent. Mice were allowed to freely explore for 10 minutes. Optogenetic stimulation was delivered as described above.

#### Small Box

This session was performed in a conditioning chamber different from the one that was used for Fear Conditioning Tagging (7 in W x 7 in D x 12 in H; Coulbourn Instruments). We taped laminated sheets with a vertical bar design to the walls of the chamber and placed a solid opaque plastic insert on the bottom of the chamber. We additionally introduced an almond scent in the room. Mice were allowed to freely explore for 10 minutes. Optogenetic stimulation was delivered as described above.

#### Natural Recall

Mice dedicated to the Natural Recall group were subjected to CFC as previously described (see Fear Conditioning and Tagging). However, these mice were not surgerized and therefore did not have any engram tagging. They were placed back in the original chamber in which they received CFC 24 hours later. The size of the chamber was similar to that of the Small Box condition (7 in W x 7 in D x 12 in H; Coulbourn Instruments) but with no alterations to walls, scent, or flooring.

### Behavioral Analysis

Video data from the Behavioral Assays were taken using GoPro cameras and analyzed using video tracking software (ANY-Maze). The total time spent in the center and total time freezing were automatically quantified and binned into two-minute intervals corresponding to the light-epoch.

### Immunohistochemistry

All mice were transcardially perfused 90-minutes after the first bin of optogenetic stimulation with 4°C phosphate buffered saline (PBS) followed by 4% paraformaldehyde (PFA) in PBS.

All brains for slice immunohistochemistry were stored in PFA for 48 hours and subsequently transferred to 30% sucrose solution one day prior to slicing. Brains were serial sectioned in 50 μm increments using a vibratome (Leica, VT100S) and collected in cold PBS. We collected slices containing the DG and the LHA as needed. All slices were incubated for two hours at room temperature in a 1x PBS + 2% Triton (PBST) and 5% normal goat serum (NGS) on an orbital shaker (Amazon). Slices were then transferred to wells containing a primary antibody solution (1:1000 rabbit anti-cFos [SySy], 1:5000 chicken anti-GFP [Invitrogen]) and were left to incubate for 48 hours on an orbital shaker at 4°C. Slices were washed with PBST for 40 minutes (20 mins x2) followed by incubation with a secondary antibody solution (1:200 Alexa 555 anti-rabbit [Invitrogen]; 1:200 Alexa 488 anti-chicken [Invitrogen]). After incubation, slices were washed once more as previously described and mounted onto microscope slides (VWR International, LLC). Cell nuclei were counterstained with DAPI added to Vectashield HardSet Mounting Medium on a coverslip and were left to dry overnight.

### Confocal Microscopy and Cell Quantification

To confirm virus expression, we acquired images using a LSM-800 confocal microscope with a 20x objective lens (Zeiss, Germany). Each image of the region of interest (ROI) contained 20 slices in a z-stack with a step size of 1.54 μm. We additionally set a series of tiles with a 10% overlap to create a single image of the ROI. Images were captured either manually with no focus strategy or were automated using the Software Autofocus feature in Zen Blue (version 2.3) to detect the most intense fluorescent pixels within the defined z-stack. All DAPI and cFos cells were quantified using a machine learning approach (Berg et al., 2019).

### LifeCanvas Technologies

Brains for network analyses were stored in PFA for 24 hours after perfusion and extraction. They were then stored in 0.02% sodium azide solution before being sent to LifeCanvas Technologies (Cambridge, MA) for brain-wide cFos detection.

Once there, brains undergo a series of preservation and clearing steps using SHIELD (Park et al., 2019) and SmartClear Pro technology (Kim et al., 2015), respectively. The samples are then washed and prepped for organ-scale immunolabeling using SmartLabel reagents (Yun et al., 2019). Samples are batch labeled in 5 μg goat anti-GFP and 3.5 μg rabbit anti-cFos per brain using SmartBatch+ and are left to incubate for roughly 18 hours. Then, samples undergo a series of washes and fixation steps over subsequent days before being incubated in secondary solutions (Yun et al., 2019). Finally, brains are mounted in agarose + EasyIndex solution for image preparation.

Brain-wide images are acquired using a SmartSPIM microscope equipped with a 3.6x objective with a 1.8 μm x 1.8 μm pixel size and a z-step size of 4 μm. The axial resolution of the images is < 4.0 μm. Tile correcting and destripping are also applied as described in Swaney *et al*. 2019. The samples are imaged using three channels: 488 nm (Autofluorescence/NeuN), 561 nm (GFP), and 642 nm (cFos).

The Autofluorescence channel is used to align the images to the Allen Brain Atlas (Allen Institute: https://portal.brain-map.org/). LifeCanvas technologies carries out this alignment process in two phases. The first phase is an automated process that samples 10-20 atlas-aligned reference samples for each brain sample using a variety of SimpleElastix warping algorithms. An average alignment was computed for all other intermediate images. To confirm the efficacy of the alignment algorithm, the second phase uses a custom Neuroglancer interface (Nuggt: https://github.com/chunglabmit/nuggt) for manual confirmation of the automated alignment algorithm. The researcher would adjust correspondence points between the Atlas and the sample image as needed to ensure more rigid alignment.

Once the images are aligned, cell populations were then mapped onto the Atlas for region-specific quantification. LifeCanvas Technologies developed a custom convolutional neural network using the Tensorflow python package (Google). Cell-detection was performed by two networks in sequence. A fully-convolutional detection network (https://arxiv.org/abs/1605.06211v1) based on a U-Net architecture (https://arxiv.org/abs/1505.04597v1) was used to find possible positive locations. Second, a convolutional network using a ResNet architecture (https://arxiv.org/abs/1512.03385v1) was used to classify each location as positive or negative. Once the cells were aligned and quantified, cFos data was aggregated into .csv files and sent back to the Ramirez Group for further analyses as described below.

### Network Generation and Analysis

Networks and all subsequent analysis thereof were generated using custom python (ver. 3.9.12) scripts built upon networkx (ver. 2.8.4), scipy (ver. 1.8.0), matplotlib (ver. 3.5.2), numpy (ver. 1.22.0), pandas (ver. 1.4.3), seaborn (ver. 0.11.2), statsmodels (ver. 0.13.2), sklearn (ver. 1.2.1), markov_clustering (ver. 0.0.06), and the brain connectivity toolbox (ver. 0.6.0). For network creation, we first finalized the number of regions of interest (ROIs) from the datasheets obtained from LifeCanvas Technologies. We discarded any region that was a layer of a ROI (e.g., Layer 2/3 of Motor Cortex) or an entry that was inclusive of several distinct ROIs (e.g., Isocortex). Additionally, we eliminated registered fiber tracts and ventricular systems. After extraneous entry elimination, we averaged hemispheric entries to obtain a bilateral density value for each ROI for each animal. From there, inter-region Spearman correlation values were calculated from these density metrics across all animals in the respective experimental condition. Thus, the networks were created such that the regions of interest were nodes, while the RS values between them were our weighted edges. Before network creation however, we chose percentile values to threshold our edges to ensure equally dense network creation between groups (e.g., a 25th percentile network removes all edges not in the top 25th percentile of edges by the largest values of RS). Intra-network communities were identified by the Louvain clustering algorithm (Blondel *et al*. 2008).

### Statistical Analysis

The sampling size for each experimental group was determined based on previous studies and are reported in figure captions. All statistics for behavioral experiments were performed using both Python and GraphPad Prism Version 9.2. Data is presented in the figures as Mean ± SEM. Behavioral data was binned at 2-minute intervals that corresponded to light-epochs (ON vs. OFF) and a repeated-measures one-way analysis of variance (ANOVA) was used to identify differences in behavior across light-epochs. Follow-up statistical analyses (two-way ANOVA, Welch’s t-test) and post-hoc analyses (Tukey’s HSD) were conducted as appropriate. Network statistical analyses were performed using Python 3.9.12. All statistical tests assumed an alpha level of 0.05. Statistical tests are reported in each figure legend with (*) = p < 0.05, (**) = p < 0.01, (***) = p < 0.001 and (****) = p < 0.0001.

## Code and Data Availability

All code is freely available in the following GitHub repository: https://github.com/rsenne/network_analysis. All data will be made available upon reasonable request.

## ACKNOWLEDGMENTS

This work was supported by a Ludwig Family Foundation, an NIH Early Independence Award (DP5 OD023106-01), an NIH Transformative R01 Award, a Young Investigator Grant from the Brain and Behavior Research Foundation, the McKnight Foundation Memory and Cognitive Disorders award, the Pew Scholars Program in the Biomedical Sciences, the Chan Zuckerberg Foundation, the Air Force Office of Scientific Research (FA9550-21-1-0310), the Neurophotonics Center at Boston University, and the Center for Systems Neuroscience and Neurophotonics Center at Boston University.

---

| **Brain Region** | **Abbreviation** | **Allen Area** |
| --- | --- | --- |
| Anterior amygdalar area | AAA | Striatum |
| Anterior cingulate area | ACA | Cortical plate |
| Nucleus accumbens | ACB | Striatum |
| Anterior hypothalamic nucleus | AHN | Hypothalamus |
| Agranular insular area | AI | Cortical plate |
| Anterior tegmental nucleus | AT | Midbrain |
| Auditory areas | AUD | Cortical plate |
| Anteroventral preoptic nucleus | AVP | Hypothalamus |
| Anteroventral periventricular nucleus | AVPV | Hypothalamus |
| Bed nucleus of the anterior commissure | BAC | Pallidum |
| Basolateral amygdala | BLA | Cortical subplate |
| Basomedial amygdala | BMA | Cortical subplate |
| Bed nuclei of the stria terminalis | BST | Pallidum |
| Field CA1 | CA1 | Cortical plate |
| Field CA2 | CA2 | Cortical plate |
| Field CA3 | CA3 | Cortical plate |
| Central amygdalar nucleus | CEA | Cortical subplate |
| Central lateral nucleus of the thalamus | CL | Thalamus |
| Clastrum | CLA | Cortical subplate |
| Central linear nucleus raphe | CLI | Midbrain |
| Central medial nucleus of the thalamus | CM | Thalamus |
| Caudoputamen | CP | Striatum |
| Superior central nucleus raphe | CS | Pons |
| Cuneiform nucleus | CUN | Midbrain |
| Dentate Gyrus | DG | Cortical plate |
| Dorsomedial nucleus of the hypothalamus | DMH | Hypothalamus |
| Denate nucleus | DN | Cerebellum |
| Dorsal nucleus raphe | DR | Midbrain |
| Ectorhinal area | ECT | Cortical plate |
| Entorhinal area | ENT | Cortical plate |
| Endopiriform nucleus | EP | Cortical subplate |
| Edinger-Westphal nucleus | EW | Midbrain |
| Fasciola cinerea | FC | Cortical plate |
| Fastigial nucleus | FN | Cerebellum |
| Fundus of striatum | FS | Striatum |
| Globus pallidus, external segment | GPe | Pallidum |
| Globus pallidus, internal segment | GPi | Pallidum |
| Gustatory areas | GU | Cortical plate |
| Hemispheric regions | HEM | Cerebellum |
| Intercalated amygdalar nucleus | IA | Striatum |
| Inferior colliculus | IC | Midbrain |
| Interfascicular nucleus raphe | IF | Midbrain |
| Induseum griseum | IG | Cortical plate |
| Infralimbic area | ILA | Cortical plate |
| Interposed nucleus | IP | Cerebellum |
| Interpeduncular nucleus | IPN | Midbrain |
| Lateral amygdalar nucleus | LA | Cortical subplate |
| Locus ceruleus | LC | Pons |
| Lateral dorsal nucleus of thalamus | LD | Thalamus |
| Lateral habenula | LH | Thalamus |
| Lateral hypothalamic area | LHA | Hypothalamus |
| Lateral preoptic area | LPO | Hypothalamus |
| Lateral septal nucleus | LS | Striatum |
| Magnocellular nucleus | MA | Pallidum |
| Medial accessory oculomotor nucleus | MA3 | Midbrain |
| Mammillary body | MBO | Hypothalamus |
| Mediodorsal nucleus of thalamus | MD | Thalamus |
| Median eminence | ME | Hypothalamus |
| Medial amygdalar nucleus | MEA | Cortical subplate |
| Median preoptic nucleus | MEPO | Hypothalamus |
| Midbrain trigeminal nucleus | MEV | Midbrain |
| Medial habenula | MH | Thalamus |
| Somatomotor areas | MO | Cortical plate |
| Medial preoptic nucleus | MPN | Hypothalamus |
| Medial preoptic area | MPO | Hypothalamus |
| Midbrain reticular nucleus | MRN | Midbrain |
| Medial septal nucleus | MS | Pallidum |
| Medulla, motor related | MY-mot | Medulla |
| Medulla, behavioral state related | MY-sat | Medulla |
| Medulla, sensory related | MY-sen | Medulla |
| Nucleus of the brachium of the inferior colliculus | NB | Midbrain |
| Diagonal band nucleus | NDB | Pallidum |
| Olfactory areas | OLF | Cortical plate |
| Orbital area | ORB | Cortical plate |
| Olfactory tubercle | OT | Striatum |
| Vascular organ and of the lamina terminalis | OV | Hypothalamus |
| Pons, motor related | P-mot | Pons |
| Pons, behavioral state related | P-sat | Pons |
| Pons, sensory related | P-sen | Pons |
| Posterior amygdalar nucleus | PA | Cortical subplate |
| Periaqueductal gray | PAG | Midbrain |
| Parasubiculum | PAR | Cortical plate |
| Parabrachial nucleus | PB | Pons |
| Parabigeminal nucleus | PBG | Midbrain |
| Paracentral nucleus | PCN | Thalamus |
| Posterodorsal preoptic nucleus | PD | Hypothalamus |
| Perirhinal area | PERI | Cortical plate |
| Parafascicular nucleus | PF | Thalamus |
| Posterior hypothalamic nucleus | PH | Hypothalamus |
| Prelimbic area | PL | Cortical plate |
| Dorsal premammillary nucleus | PMd | Hypothalamus |
| Ventral premammillary nucleus | PMv | Hypothalamus |
| Paranigral nucleus | PN | Midbrain |
| Postsubiculum | POST | Cortical plate |
| Peripeduncular nucleus | PP | Thalamus |
| Pedunculopontine nucleus | PPN | Midbrain |
| Perireunensis nucleus | PR | Thalamus |
| Presubiculum | PRE | Cortical plate |
| Pretectal region | PRT | Midbrain |
| Parastrial nucleus | PS | Hypothalamus |
| Preparasubthalamic nucleus | PST | Hypothalamus |
| Parasubthalamic nucleus | PSTN | Hypothalamus |
| Posterior parietal association areas | PTLp | Cortical plate |
| Paraventricular nucleus of the thalamus | PVT | Thalamus |
| Periventricular zone | PVZ | Hypothalamus |
| Perifornical nucleus | PeF | Hypothalamus |
| Prosubiculum | ProS | Cortical plate |
| Retrochiasmatic area | RCH | Hypothalamus |
| Nucleus of reuniens | RE | Thalamus |
| Rhomboid nucleus | RH | Thalamus |
| Rostral linear nucleus raphe | RL | Midbrain |
| Red nucleus | RN | Midbrain |
| Midbrain reticular nucleus, retrorubral area | RR | Midbrain |
| Retrosplenial area | RSP | Cortical plate |
| Reticular nucleus of the thalamus | RT | Thalamus |
| Nucleus sagulum | SAG | Midbrain |
| Subparaventricular zone | SBPV | Hypothalamus |
| Suprachiasmatic nucleus | SCH | Hypothalamus |
| Superior colliculus, motor related | SCm | Midbrain |
| Superior colliculus, sensory related | SCs | Midbrain |
| Septofimbrial nucleus | SF | Striatum |
| Subfornical organ | SFO | Hypothalamus |
| Septohippocampal nucleus | SH | Striatum |
| Substantia innominata | SI | Pallidum |
| Substantia nigra, compact part | SNC | Midbrain |
| Substantia nigra, reticular part | SNr | Midbrain |
| Subparafascicular area | SPA | Thalamus |
| Subparafascicular nucleus | SPF | Thalamus |
| Somatosensory areas | SS | Cortical plate |
| Subthalamic nucleus | STN | Hypothalamus |
| Subiculum | SUB | Cortical plate |
| Temporal association areas | TEa | Cortical plate |
| Triangular nucleus of septum | TRS | Pallidum |
| Tuberal nucleus | TU | Hypothalamus |
| Ventral anterior-lateral complex of the thalamus | VAL | Thalamus |
| Vermal regions | VERM | Cerebellum |
| Visual areas | VIS | Cortical plate |
| Visceral area | VISC | Cortical plate |
| Ventrolateral preoptic nucleus | VLPO | Hypothalamus |
| Ventral medial nucleus of the thalamus | VM | Thalamus |
| Ventromedial hypothalamic nucleus | VMH | Hypothalamus |
| Ventromedial preoptic nucleus | VMPO | Hypothalamus |
| Ventral posterior complex of the thalamus | VP | Thalamus |
| Ventral tegmental area | VTa | Midbrain |
| Ventral tegmental nucleus | VTN | Midbrain |
| Vestibulocerebellar nucleus | VeCB | Cerebellum |
| Zona incerta | ZI | Hypothalamus |

## REFERENCES

Amano, T., Duvarci, S., Popa, D., Pare, D., 2011. The Fear Circuit Revisited: Contributions of the Basal Amygdala Nuclei to Conditioned Fear. J. Neurosci. 31, 15481–15489. https://doi.org/10.1523/JNEUROSCI.3410-11.2011

Benjamini, Y., Hochberg, Y., 1995. Controlling the False Discovery Rate: A Practical and Powerful Approach to Multiple Testing. J. R. Stat. Soc. Ser. B Methodol. 57, 289–300. https://doi.org/10.1111/j.2517-6161.1995.tb02031.x

Berg, S., Kutra, D., Kroeger, T., Straehle, C.N., Kausler, B.X., Haubold, C., Schiegg, M., Ales, J., Beier, T., Rudy, M., Eren, K., Cervantes, J.I., Xu, B., Beuttenmueller, F., Wolny, A., Zhang, C., Koethe, U., Hamprecht, F.A., Kreshuk, A., 2019. ilastik: interactive machine learning for (bio)image analysis. Nat. Methods 16, 1226–1232. https://doi.org/10.1038/s41592-019-0582-9

Blanchard, D.C., Blanchard, R.J., 2008. Chapter 2.4 Defensive behaviors, fear, and anxiety, in: Blanchard, R.J., Blanchard, D.C., Griebel, G., Nutt, D. (Eds.), Handbook of Behavioral Neuroscience, Handbook of Anxiety and Fear. Elsevier, pp. 63–79. https://doi.org/10.1016/S1569-7339(07)00005-7

Blanchard, R.J., Blanchard, D.C., 1989. Attack and defense in rodents as ethoexperimental models for the study of emotion. Prog. Neuropsychopharmacol. Biol. Psychiatry 13, S3–S14. https://doi.org/10.1016/0278-5846(89)90105-X

Blondel, V.D., Guillaume, J.-L., Lambiotte, R., Lefebvre, E., 2008. Fast unfolding of communities in large networks. J. Stat. Mech. Theory Exp. 2008, P10008. https://doi.org/10.1088/1742-5468/2008/10/P10008

Bonnavion, P., Jackson, A.C., Carter, M.E., de Lecea, L., 2015. Antagonistic interplay between hypocretin and leptin in the lateral hypothalamus regulates stress responses. Nat. Commun. 6, 6266. https://doi.org/10.1038/ncomms7266

Chen, B.K., Murawski, N.J., Cincotta, C., McKissick, O., Finkelstein, A., Hamidi, A.B., Merfeld, E., Doucette, E., Grella, S.L., Shpokayte, M., Zaki, Y., Fortin, A., Ramirez, S., 2019. Artificially Enhancing and Suppressing Hippocampus-Mediated Memories. Curr. Biol. 29, 1885–1894.e4. https://doi.org/10.1016/j.cub.2019.04.065

Cho, J.-H., Rendall, S.D., Gray, J.M., 2017. Brain-wide maps of *Fos* expression during fear learning and recall. Learn. Mem. 24, 169–181. https://doi.org/10.1101/lm.044446.116

Coelho, C.A.O., Ferreira, T.L., Kramer-Soares, J.C., Sato, J.R., Oliveira, M.G.M., 2018. Network supporting contextual fear learning after dorsal hippocampal damage has increased dependence on retrosplenial cortex. PLOS Comput. Biol. 14, e1006207. https://doi.org/10.1371/journal.pcbi.1006207

Colom-Lapetina, J., Li, A.J., Pelegrina-Perez, T.C., Shansky, R.M., 2019. Behavioral Diversity Across Classic Rodent Models Is Sex-Dependent. Front. Behav. Neurosci. 13, 45. https://doi.org/10.3389/fnbeh.2019.00045

Concetti, C., Bracey, E.F., Peleg-Raibstein, D., Burdakov, D., 2020. Control of fear extinction by hypothalamic melanin-concentrating hormone–expressing neurons. Proc. Natl. Acad. Sci. 117, 22514–22521. https://doi.org/10.1073/pnas.2007993117

Dean, K.M., Roudot, P., Welf, E.S., Danuser, G., Fiolka, R., 2015. Deconvolution-free Subcellular Imaging with Axially Swept Light Sheet Microscopy. Biophys. J. 108, 2807–2815. https://doi.org/10.1016/j.bpj.2015.05.013

Deng, H., Xiao, X., Wang, Z., 2016. Periaqueductal Gray Neuronal Activities Underlie Different Aspects of Defensive Behaviors. J. Neurosci. 36, 7580–7588. https://doi.org/10.1523/JNEUROSCI.4425-15.2016

Do-Monte, F.H., Quiñones-Laracuente, K., Quirk, G.J., 2015. A temporal shift in the circuits mediating retrieval of fear memory. Nature 519, 460–463. https://doi.org/10.1038/nature14030

Fadok, J.P., Krabbe, S., Markovic, M., Courtin, J., Xu, C., Massi, L., Botta, P., Bylund, K., Müller, C., Kovacevic, A., Tovote, P., Lüthi, A., 2017. A competitive inhibitory circuit for selection of active and passive fear responses. Nature 542, 96–100. https://doi.org/10.1038/nature21047

Fanselow, M.S., 1994. Neural organization of the defensive behavior system responsible for fear. Psychon. Bull. Rev. 1, 429–438. https://doi.org/10.3758/BF03210947

Fanselow, M.S., Dong, H.-W., 2010. Are the Dorsal and Ventral Hippocampus Functionally Distinct Structures? Neuron 65, 7–19.https://doi.org/10.1016/j.neuron.2009.11.031

Fendt, M., Fanselow, M.S., 1999. The neuroanatomical and neurochemical basis of conditioned fear. Neurosci. Biobehav. Rev. 23, 743–760. https://doi.org/10.1016/S0149-7634(99)00016-0

Garrison, K.A., Scheinost, D., Finn, E.S., Shen, X., Constable, R.T., 2015. The (in)stability of functional brain network measures across thresholds. NeuroImage 118, 651–661. https://doi.org/10.1016/j.neuroimage.2015.05.046

Glickman, M.E., Rao, S.R., Schultz, M.R., 2014. False discovery rate control is a recommended alternative to Bonferroni-type adjustments in health studies. J. Clin. Epidemiol. 67, 850–857. https://doi.org/10.1016/j.jclinepi.2014.03.012

Gross, C.T., Canteras, N.S., 2012. The many paths to fear. Nat. Rev. Neurosci. 13, 651–658. https://doi.org/10.1038/nrn3301

Grossen, N.E., Kelley, M.J., 1972. Species-specific behavior and acquisition of avoidance behavior in rats. J. Comp. Physiol. Psychol. 81, 307–310. https://doi.org/10.1037/h0033536

Gruene, T.M., Flick, K., Stefano, A., Shea, S.D., Shansky, R.M., 2015. Sexually divergent expression of active and passive conditioned fear responses in rats. eLife 4, e11352. https://doi.org/10.7554/eLife.11352

Guimerà, R., Nunes Amaral, L.A., 2005. Functional cartography of complex metabolic networks. Nature 433, 895–900. https://doi.org/10.1038/nature03288

Herry, C., Johansen, J.P., 2014. Encoding of fear learning and memory in distributed neuronal circuits. Nat. Neurosci. 17, 1644–1654. https://doi.org/10.1038/nn.3869

Jardim, M.C., Guimarães, F.S., 2004. Role of glutamate ionotropic receptors in the dorsomedial hypothalamic nucleus on anxiety and locomotor behavior. Pharmacol. Biochem. Behav. 79, 541–546. https://doi.org/10.1016/j.pbb.2004.09.005

Jimenez, J.C., Su, K., Goldberg, A.R., Luna, V.M., Biane, J.S., Ordek, G., Zhou, P., Ong, S.K., Wright, M.A., Zweifel, L., Paninski, L., Hen, R., Kheirbek, M.A., 2018. Anxiety Cells in a Hippocampal-Hypothalamic Circuit. Neuron 97, 670–683.e6. https://doi.org/10.1016/j.neuron.2018.01.016

Josselyn, S.A., Tonegawa, S., 2020. Memory engrams: Recalling the past and imagining the future. Science 367, eaaw4325. https://doi.org/10.1126/science.aaw4325

Kalueff, A.V., Stewart, A.M., Song, C., Berridge, K.C., Graybiel, A.M., Fentress, J.C., 2016. Neurobiology of rodent self-grooming and its value for translational neuroscience. Nat. Rev. Neurosci. 17, 45–59. https://doi.org/10.1038/nrn.2015.8

Kim, J.J., Rison, R.A., Fanselow, M.S., 1993. Effects of amygdala, hippocampus, and periaqueductal gray lesions on short-and long-term contextual fear. Behav. Neurosci. 107, 1093–1098. https://doi.org/10.1037/0735-7044.107.6.1093

Kim, S.-Y., Adhikari, A., Lee, S.Y., Marshel, J.H., Kim, C.K., Mallory, C.S., Lo, M., Pak, S., Mattis, J., Lim, B.K., Malenka, R.C., Warden, M.R., Neve, R., Tye, K.M., Deisseroth, K., 2013. Diverging neural pathways assemble a behavioural state from separable features in anxiety. Nature 496, 219–223. https://doi.org/10.1038/nature12018

Kim, S.-Y., Cho, J.H., Murray, E., Bakh, N., Choi, H., Ohn, K., Ruelas, L., Hubbert, A., McCue, M., Vassallo, S.L., Keller, P.J., Chung, K., 2015. Stochastic electrotransport selectively enhances the transport of highly electromobile molecules. Proc. Natl. Acad. Sci. 112. https://doi.org/10.1073/pnas.1510133112

Lin, D., Boyle, M.P., Dollar, P., Lee, H., Lein, E.S., Perona, P., Anderson, D.J., 2011. Functional identification of an aggression locus in the mouse hypothalamus. Nature 470, 221–226. https://doi.org/10.1038/nature09736

Liu, X., Ramirez, S., Pang, P.T., Puryear, C.B., Govindarajan, A., Deisseroth, K., Tonegawa, S., 2012.Optogenetic stimulation of a hippocampal engram activates fear memory recall. Nature 484, 381–385. https://doi.org/10.1038/nature11028

Ma, J., du Hoffmann, J., Kindel, M., Beas, B.S., Chudasama, Y., Penzo, M.A., 2021. Divergent projections of the paraventricular nucleus of the thalamus mediate the selection of passive and active defensive behaviors. Nat. Neurosci. 24, 1429–1440. https://doi.org/10.1038/s41593-021-00912-7

Mangieri, L.R., Jiang, Z., Lu, Y., Xu, Yuanzhong, Cassidy, R.M., Justice, N., Xu, Yong, Arenkiel, B.R., Tong, Q., 2019. Defensive Behaviors Driven by a Hypothalamic-Ventral Midbrain Circuit. eneuro 6, ENEURO.0156-19.2019. https://doi.org/10.1523/ENEURO.0156-19.2019

Maren, S., Phan, K.L., Liberzon, I., 2013. The contextual brain: implications for fear conditioning, extinction and psychopathology. Nat. Rev. Neurosci. 14, 417–428. https://doi.org/10.1038/nrn3492

McInnes, L., Healy, J., Melville, J., 2020. UMAP: Uniform Manifold Approximation and Projection for Dimension Reduction.

Molewijk, H.E., 1995. The ambivalent behaviour “stretched approach posture” in the rat as a paradigm to characterize anxiolyUc drugs.

Park, Y.-G., Sohn, C.H., Chen, R., McCue, M., Yun, D.H., Drummond, G.T., Ku, T., Evans, N.B., Oak, H.C., Trieu, W., Choi, H., Jin, X., Lilascharoen, V., Wang, J., Truttmann, M.C., Qi, H.W., Ploegh, H.L., Golub, T.R., Chen, S.-C., Frosch, M.P., Kulik, H.J., Lim, B.K., Chung, K., 2019. Protection of tissue physicochemical properties using polyfunctional crosslinkers. Nat. Biotechnol. 37, 73–83. https://doi.org/10.1038/nbt.4281

Penzo, M.A., Robert, V., Tucciarone, J., De Bundel, D., Wang, M., Van Aelst, L., Darvas, M., Parada, L.F., Palmiter, R.D., He, M., Huang, Z.J., Li, B., 2015. The paraventricular thalamus controls a central amygdala fear circuit. Nature 519, 455–459. https://doi.org/10.1038/nature13978

Pobbe, R.L.H., Zangrossi, H., 2010. The lateral habenula regulates defensive behaviors through changes in 5-HT-mediated neurotransmission in the dorsal periaqueductal gray matter. Neurosci. Lett. 479, 87–91. https://doi.org/10.1016/j.neulet.2010.05.021

Ramirez, S., Liu, X., Lin, P.-A., Suh, J., Pignatelli, M., Redondo, R.L., Ryan, T.J., Tonegawa, S., 2013. Creating a False Memory in the Hippocampus. Science 341, 387–391. https://doi.org/10.1126/science.1239073

Ramirez, S., Liu, X., MacDonald, C.J., Moffa, A., Zhou, J., Redondo, R.L., Tonegawa, S., 2015. Activating positive memory engrams suppresses depression-like behaviour. Nature 522, 335–339. https://doi.org/10.1038/nature14514

Redondo, R.L., Kim, J., Arons, A.L., Ramirez, S., Liu, X., Tonegawa, S., 2014. Bidirectional switch of the valence associated with a hippocampal contextual memory engram. Nature 513, 426–430. https://doi.org/10.1038/nature13725

Reijmers, L.G., Perkins, B.L., Matsuo, N., Mayford, M., 2007. Localization of a Stable Neural Correlate of Associative Memory. Science 317, 1230–1233. https://doi.org/10.1126/science.1143839

Rosen, J.B., Pagani, J.H., Rolla, K.L.G., Davis, C., 2008. Analysis of behavioral constraints and the neuroanatomy of fear to the predator odor trimethylthiazoline: A model for animal phobias. Neurosci. Biobehav. Rev. 32, 1267–1276. https://doi.org/10.1016/j.neubiorev.2008.05.006

Roy, D.S., Arons, A., Mitchell, T.I., Pignatelli, M., Ryan, T.J., Tonegawa, S., 2016. Memory retrieval by activating engram cells in mouse models of early Alzheimer’s disease. Nature 531, 508–512. https://doi.org/10.1038/nature17172

Roy, D.S., Park, Y.-G., Kim, M.E., Zhang, Y., Ogawa, S.K., DiNapoli, N., Gu, X., Cho, J.H., Choi, H., Kamentsky, L., Martin, J., Mosto, O., Aida, T., Chung, K., Tonegawa, S., 2022. Brain-wide mapping reveals that engrams for a single memory are distributed across multiple brain regions. Nat. Commun. 13, 1799. https://doi.org/10.1038/s41467-022-29384-4

Rubinov, M., Sporns, O., 2011. Weight-conserving characterization of complex functional brain networks. NeuroImage 56, 2068–2079. https://doi.org/10.1016/j.neuroimage.2011.03.069

Ryan, T.J., Roy, D.S., Pignatelli, M., Arons, A., Tonegawa, S., 2015. Engram cells retain memory under retrograde amnesia. Science 348, 1007–1013. https://doi.org/10.1126/science.aaa5542

Salay, L.D., Ishiko, N., Huberman, A.D., 2018. A midline thalamic circuit determines reactions to visual threat. Nature 557, 183–189. https://doi.org/10.1038/s41586-018-0078-2

Scoviille, W.B., Milner, B., 1957. LOSS OF RECENT MEMORY AFTER BILATERAL HIPPOCAMPAL LESIONS.

Shansky, R.M., Murphy, A.Z., 2021. Considering sex as a biological variable will require a global shift in science culture. Nat. Neurosci. 24, 457–464. https://doi.org/10.1038/s41593-021-00806-8

Silva, B.A., Gross, C.T., Gräff, J., 2016. The neural circuits of innate fear: detection, integration, action, and memorization. Learn. Mem. 23, 544–555. https://doi.org/10.1101/lm.042812.116

Song, C., Berridge, K.C., Kalueff, A.V., 2016. “Stressing” rodent self-grooming for neuroscience research. Nat. Rev. Neurosci. 17, 591–591. https://doi.org/10.1038/nrn.2016.103

Soria-Gómez, E., Busquets-Garcia, A., Hu, F., Mehidi, A., Cannich, A., Roux, L., Louit, I., Alonso, L., Wiesner, T., Georges, F., Verrier, D., Vincent, P., Ferreira, G., Luo, M., Marsicano, G., 2015. Habenular CB1 Receptors Control the Expression of Aversive Memories. Neuron 88, 306–313. https://doi.org/10.1016/j.neuron.2015.08.035

Stamatakis, A.M., Stuber, G.D., 2012. Activation of lateral habenula inputs to the ventral midbrain promotes behavioral avoidance. Nat. Neurosci. 15, 1105–1107. https://doi.org/10.1038/nn.3145

Swaney, J., Kamentsky, L., Evans, N.B., Xie, K., Park, Y.-G., Drummond, G., Yun, D.H., Chung, K., 2019. Scalable image processing techniques for quantitative analysis of volumetric biological images from light-sheet microscopy (preprint). Bioengineering. https://doi.org/10.1101/576595

Tovote, P., Esposito, M.S., Botta, P., Chaudun, F., Fadok, J.P., Markovic, M., Wolff, S.B.E., Ramakrishnan, C., Fenno, L., Deisseroth, K., Herry, C., Arber, S., Lüthi, A., 2016. Midbrain circuits for defensive behaviour. Nature 534, 206–212. https://doi.org/10.1038/nature17996

Tovote, P., Fadok, J.P., Lüthi, A., 2015. Neuronal circuits for fear and anxiety. Nat. Rev. Neurosci. 16, 317–331. https://doi.org/10.1038/nrn3945

Vale, R., Evans, D.A., Branco, T., 2017. Rapid Spatial Learning Controls Instinctive Defensive Behavior in Mice. Curr. Biol. 27, 1342–1349. https://doi.org/10.1016/j.cub.2017.03.031

van den Heuvel, M.P., Mandl, R.C.W., Stam, C.J., Kahn, R.S., Hulshoff Pol, H.E., 2010. Aberrant Frontal and Temporal Complex Network Structure in Schizophrenia: A Graph Theoretical Analysis. J. Neurosci. 30, 15915–15926. https://doi.org/10.1523/JNEUROSCI.2874-10.2010

van den Heuvel, M.P., Sporns, O., 2013. Network hubs in the human brain. Trends Cogn. Sci. 17, 683–696. https://doi.org/10.1016/j.tics.2013.09.012

Vetere, G., Kenney, J.W., Tran, L.M., Xia, F., Steadman, P.E., Parkinson, J., Josselyn, S.A., Frankland, P.W., 2017. Chemogenetic Interrogation of a Brain-wide Fear Memory Network in Mice. Neuron 94, 363–374.e4. https://doi.org/10.1016/j.neuron.2017.03.037

Wallace, K.J., Rosen, J.B., 2000. Predator odor as an unconditioned fear stimulus in rats: Elicitation of freezing by trimethylthiazoline, a component of fox feces. Behav. Neurosci. 114, 912–922. https://doi.org/10.1037/0735-7044.114.5.912

Wang, W., Schuette, P.J., Nagai, J., Tobias, B.C., Cuccovia V. Reis, F.M., Ji, S., de Lima, M.A.X., La-Vu, M.Q., Maesta-Pereira, S., Chakerian, M., Leonard, S.J., Lin, L., Severino, A.L., Cahill, C.M., Canteras, N.S., Khakh, B.S., Kao, J.C., Adhikari, A., 2021. Coordination of escape and spatial navigation circuits orchestrates versatile flight from threats. Neuron 109, 1848–1860.e8. https://doi.org/10.1016/j.neuron.2021.03.033

Wheeler, A.L., Teixeira, C.M., Wang, A.H., Xiong, X., Kovacevic, N., Lerch, J.P., McIntosh, A.R., Parkinson, J., Frankland, P.W., 2013. Identification of a Functional Connectome for Long-Term Fear Memory in Mice. PLoS Comput. Biol. 9, e1002853. https://doi.org/10.1371/journal.pcbi.1002853

Yu, K., Garcia da Silva, P., Albeanu, D.F., Li, B., 2016. Central Amygdala Somatostatin Neurons Gate Passive and Active Defensive Behaviors. J. Neurosci. 36, 6488–6496. https://doi.org/10.1523/JNEUROSCI.4419-15.2016

Yun, D.H., Park, Y.-G., Cho, J.H., Kamentsky, L., Evans, N.B., Albanese, A., Xie, K., Swaney, J., Sohn, C.H., Tian, Y., Zhang, Q., Drummond, G., Guan, W., DiNapoli, N., Choi, H., Jung, H.-Y., Ruelas, L., Feng, G., Chung, K., 2019. Ultrafast immunostaining of organ-scale tissues for scalable proteomic phenotyping (preprint). Bioengineering. https://doi.org/10.1101/660373

Zhang, J., Tan, L., Ren, Y., Liang, J., Lin, R., Feng, Q., Zhou, J., Hu, F., Ren, J., Wei, C., Yu, T., Zhuang, Y., Bettler, B., Wang, F., Luo, M., 2016. Presynaptic Excitation via GABA B Receptors in Habenula Cholinergic Neurons Regulates Fear Memory Expression. Cell 166, 716–728. https://doi.org/10.1016/j.cell.2016.06.026

